# Fatty acid photodecarboxylase is an ancient photoenzyme responsible for hydrocarbon formation in the thylakoid membranes of algae

**DOI:** 10.1101/2020.06.23.166330

**Authors:** Solène Moulin, Audrey Beyly, Stéphanie Blangy, Bertrand Légeret, Magali Floriani, Adrien Burlacot, Damien Sorigué, Yonghua Li-Beisson, Gilles Peltier, Fred Beisson

## Abstract

Fatty acid photodecarboxylase (FAP) is one of the three enzymes that require light for their catalytic cycle (photoenzymes). FAP has been first identified in the green microalga *Chlorella variabilis* NC64A and belongs an algae-specific subgroup of the glucose-methanol-choline oxidoreductase family. While the FAP from *Chlorella* and its *Chlamydomonas reinhardtii* homolog CrFAP have demonstrated *in vitro* activity, their activity and physiological function have not been studied *in vivo*. Besides, the conservation of FAP activity beyond green microalgae remains hypothetical. Here, using a *Chlamydomonas* FAP knockout line (*fap*), we show that CrFAP is responsible for the formation of 7-heptadecene, the only hydrocarbon present in this alga. We further show that CrFAP is associated to the thylakoids and that 90% of 7-heptadecene is recovered in this cell fraction. In the *fap* mutant, photosynthesis activity was not affected under standard growth conditions but was reduced after cold acclimation. A phylogenetic analysis including sequences from Tara Ocean identified almost 200 putative FAPs and indicated that FAP was acquired early after primary endosymbiosis. Within Bikonta, FAP was kept in photosynthetic secondary endosymbiosis lineages but absent in those that lost the plastid. Characterization of recombinant FAPs from various algal genera (*Nannochloropsis, Ectocarpus, Galdieria, Chondrus*) provided experimental evidence that FAP activity is conserved in red and brown algae and is not limited to unicellular species. These results thus indicate that FAP has been conserved during evolution of most algal lineages when photosynthesis was kept and suggest that its function is linked to photosynthetic membranes.

**One sentence summary:** FAP is present in thylakoids and conserved beyond green algae.

## INTRODUCTION

Most organisms have the ability to synthesize highly hydrophobic compounds made only of carbon and hydrogen called hydrocarbons (HCs). Many HCs are isoprenoids but others like *n*-alkanes and their unsaturated analogues (*n*-alkenes) derive from fatty acids (Herman and Zhang 2016). In plants, C29-C35 *n*-alkanes are synthesized in epidermis from very-long-chain fatty acids and secreted onto the surface of aerial organs (Lee and Suh 2013). Plant *n*-alkanes are important for adaptation to the terrestrial environment because they constitute a major part of the cuticular wax layer that prevents the loss of internal water. In microorganisms, *n*-alka(e)nes are presumably mostly located in membranes and their function is more elusive. Roles in cell growth, cell division, photosynthesis and salt tolerance have been proposed for cyanobacterial *n*-alka(e)nes (Berla *et al*. 2015; Lea-Smith *et al*. 2016; Yamamori *et al*. 2018; Knoot and Pakrasi 2019). In eukaryotic microalgae, occurrence of *n*-alka(e)nes has been reported their subcellular location and physiological function remain unknown (Sorigué *et al*. 2016). Besides the elucidation of their biological roles, *n*-alkanes and *n*-alkenes have also attracted attention because of their biotechnological interest. Indeed, a bio-based alka(e)ne production would be highly desirable to replace part of petroleum-derived HCs in fuels, cosmetics, lubricants and as synthons in organic chemistry (Jetter and Kunst 2008).

A number of *n*-alka(e)ne-forming enzymes have been identified and characterized in the last decade and it is now clear that conversion of fatty acids to HCs occurs through a variety of reactions and proteins (Herman and Zhang 2016). Besides, for the same type of reaction, the biosynthetic enzymes involved are not conserved across phylogenetic groups. For instance, it has been shown in bacteria that synthesis of terminal olefins (1-alkenes) occurs through decarboxylation of a saturated long-chain fatty acid and that this reaction is catalyzed by a cytochrome P450 in *Jeotgalicoccus* spp. (Rude *et al*. 2011) and a non-heme iron oxidase in Pseudomonas (Rui *et al*. 2014). In the bacterium *Micrococcus luteus*, yet another pathway has been described, which consists in the head-to-head condensation of fatty acids to form very-long-chain *n*-alkenes with internal double bonds (Beller *et al*. 2010). In cyanobacteria, *n*-alka(e)nes are produced by two distinct pathways which are mutually exclusive (Coates *et al*. 2014). The first one forms terminal olefins and involves a type I polyketide synthase that elongates and decarboxylates a C_n_ fatty acid to yield a C_n+1_ alkene (Mendez-Perez *et al*. 2011). The second one is a two-step pathway with a fatty aldehyde intermediate and involves an acyl-ACP reductase (AAR) and an aldehyde deformylating oxygenase (ADO) (Schirmer *et al*. 2010; Li *et al*. 2012). In plants, the pathway producing the very-long-chain *n*-alkanes of the cuticular waxes is known to require the Arabidopsis homologs CER1 and CER3 and would involve an aldehyde intermediate (Bernard *et al*. 2012).

In microalgae, we have shown that C15-C17 *n*-alka(e)nes occur in *Chlorella variabilis* NC64A (named *Chlorella* from here on) and that they are synthesized through decarboxylation of long-chain fatty acids (Sorigué *et al*. 2016). A *Chlorella* protein with a fatty acid decarboxylase activity was then identified as a photoenzyme (Sorigué *et al*. 2017), a rare type of enzyme that requires light as energy (Bjorn 2015). The *Chlorella* protein was thus named fatty acid photodecarboxylase (FAP, E.C. 4.1.1.106). It is one of the three photoenzymes discovered so far, the two others being DNA photolyases and light-dependent protochlorophyllide oxidoreductase. FAP belongs to a family of flavoproteins (Sorigué *et al*. 2017), the glucose-methanol-choline (GMC) oxidoreductases, which includes a large variety of enzymes present in prokaryotic and eukaryotic organisms (Zamocky *et al*. 2004). FAP activity thus represents a new type of chemistry in the GMC oxidoreductase family (Sorigué *et al*. 2017). Molecular phylogeny has shown that *Chlorella* FAP and CrFAP belong to an algal branch of GMC oxidoreductases. However, whether FAP activity is conserved in other algal lineages beyond green algae and whether FAP is indeed responsible for *n*-alka(e)ne formation *in vivo* remains to be demonstrated. Besides, the subcellular location and role of FAP in the algal cells have not yet been investigated.

In this work, we isolate and characterize in *Chlamydomonas* reinhardtii (hereafter named *Chlamydomonas*) an insertional mutant deficient in FAP (*fap* mutant strain). We show that FAP is indeed responsible for the formation of 7-heptadecene, the only HC present in this alga. In addition, we provide evidence for a thylakoid localization of *Chlamydomonas* FAP and its alkene product. We also show that although growth and photosynthesis are not affected in the knockout under laboratory conditions, but photosynthetic efficiency is impacted under cold when light intensity varies. Finally, we build a large molecular phylogeny of GMC oxidoreductases based on TARA Ocean data and identify almost 200 new putative FAP sequences across algal lineages. Experimental evidence is provided that FAP photochemical activity is conserved in red and brown algae and is not limited to unicellular species but is also present in macroalgae.

## RESULTS

### FAP is responsible for alkene synthesis in *Chlamydomonas*

*Chlamydomonas* has been previously shown to produce 7-heptadecene (C17:1-alkene) from *cis*-vaccenic acid (Sorigué *et al*., 2016) and to have a FAP homolog that can also perform photodecarboxylation of fatty acids *in vitro* (Sorigué *et al*., 2017). Although it seemed likely that FAP proteins are indeed responsible for the synthesis of alka(e)nes produced by *Chlamydomonas* and *Chlorella*, the possibility that the alkenes are formed *in vivo* by another enzyme could not be ruled out. In order to address this issue and investigate the biological role of FAP, a *Chlamydomonas* strain mutated for FAP was isolated from the *Chlamydomonas* library project (CliP)(Li *et al*., 2016). This strain showed complete absence of FAP protein **(Fig. 1)** and was named *fap* from here on. The only HC in *Chlamydomonas* i.e. 7-heptadecene could not be detected in the *fap* mutant **(Fig. 1)**. After performing nuclear complementation using the genomic *FAP* gene under the promotor PsaD, 4 independent transformants with different expression levels of *Chlamydomonas* FAP (CrFAP) were isolated (named Cp 1 to 4). In these complemented strains, production of 7-heptadecene was clearly related to FAP amount. These results thus demonstrated that CrFAP is indeed responsible for alkene formation *in vivo* in *Chlamydomonas*.

**Figure 1:**
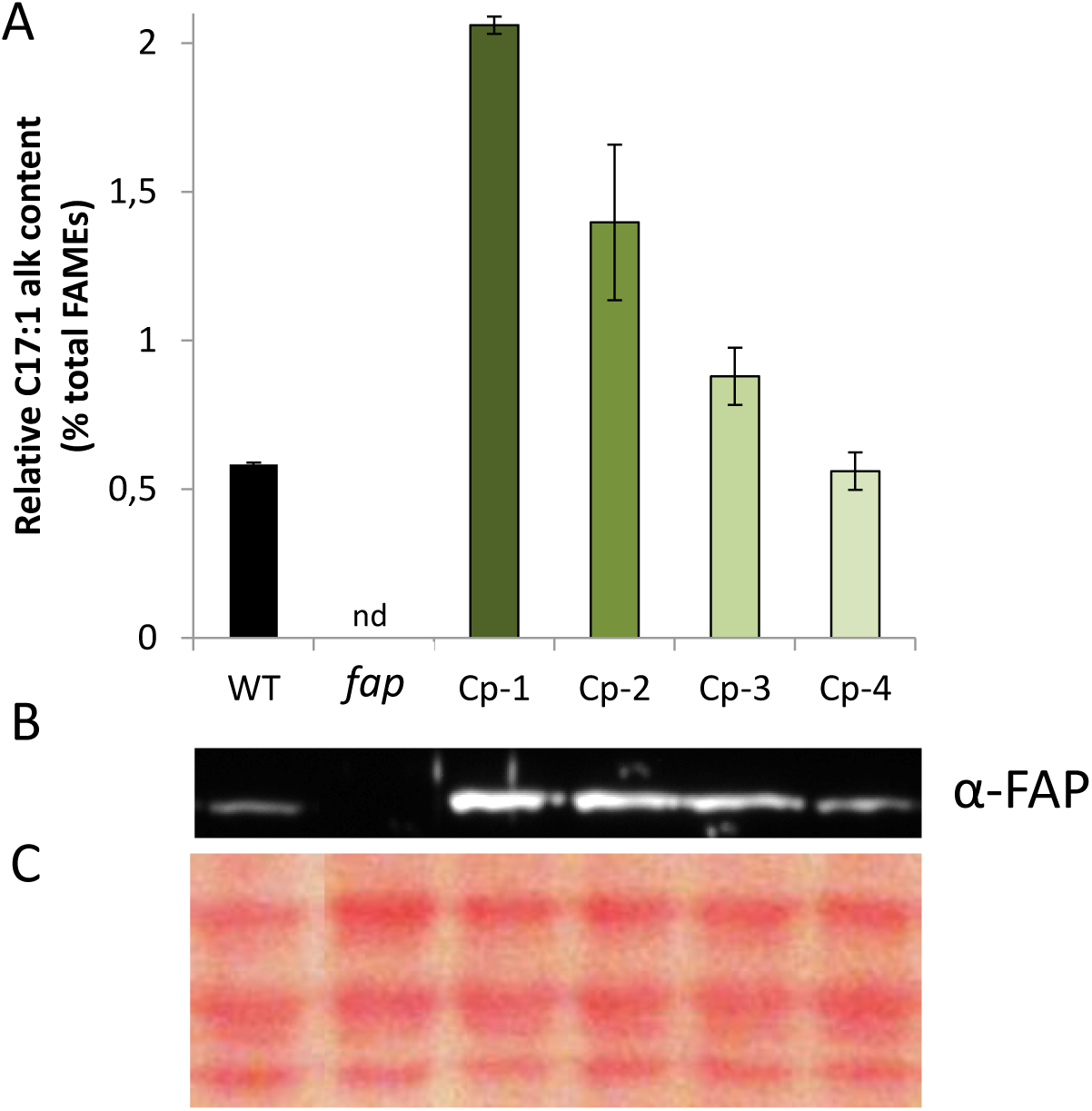
FAP level is correlated with amount of 7-heptadecene in *Chlamydomonas*. **A**, relative content in total fatty-acid derived hydrocarbons measured on whole cells; the only fatty acid-derived hydrocarbon of *Chlamydomonas* is 7-heptadecene (abbreviated as C17:1 alk). **B**, Immunoblot of total protein extract probed with anti-FAP antibody. **C**, Loading control of the immunoblot (Ponceau staining). WT: wild type strain; fap: FAP knockout; Cp 1 to 4: complemented strains; nd: not detected (strain labels in panel A correspond to lanes of panels B and C). Values are mean ± SD of n=4 independent experiments for each strain.

### FAP activity is conserved beyond green microalgae

Molecular phylogeny of GMC oxidoreductases has previously shown that CrFAP and the FAP from *Chlorella variabilis* NC64A (CvFAP) are present in a branch containing only sequences from algae (Sorigué *et al*., 2017). The term “algae” is used here in the classical sense of photosynthetic organisms that have chlorophyll *a* as their primary photosynthetic pigment and lack a sterile covering of cells around the reproductive cells (Lee, 2008). To investigate whether FAP activity has been conserved in other algal groups than green algae, genes encoding putative FAPs from selected algal lineages were cloned and expressed in *E. coli* and the bacterial HC content was analyzed. Considering the basal position of red algae, we decided to explore FAP activity in Rhodophytes selecting the microalga *Galdieria sulphuraria* and the macroalga *Chondrus crispus*. For algae deriving from secondary endosymbiosis, we also chose the microalga *Nannochloropsis gaditana* and the macroalga *Ectocarpus silicosus. E. coli* strains expressing the various FAPs all produced a range of *n*-alkanes and *n*-alkenes with different chain lengths (C15 to C17) in various proportions **(Fig. 2 and Supplemental Fig. S1)**. These results thus demonstrate that FAP activity is present in red algae, and has been conserved in algae with secondary plastids and is not limited to unicellular algae.

**Figure 2.**
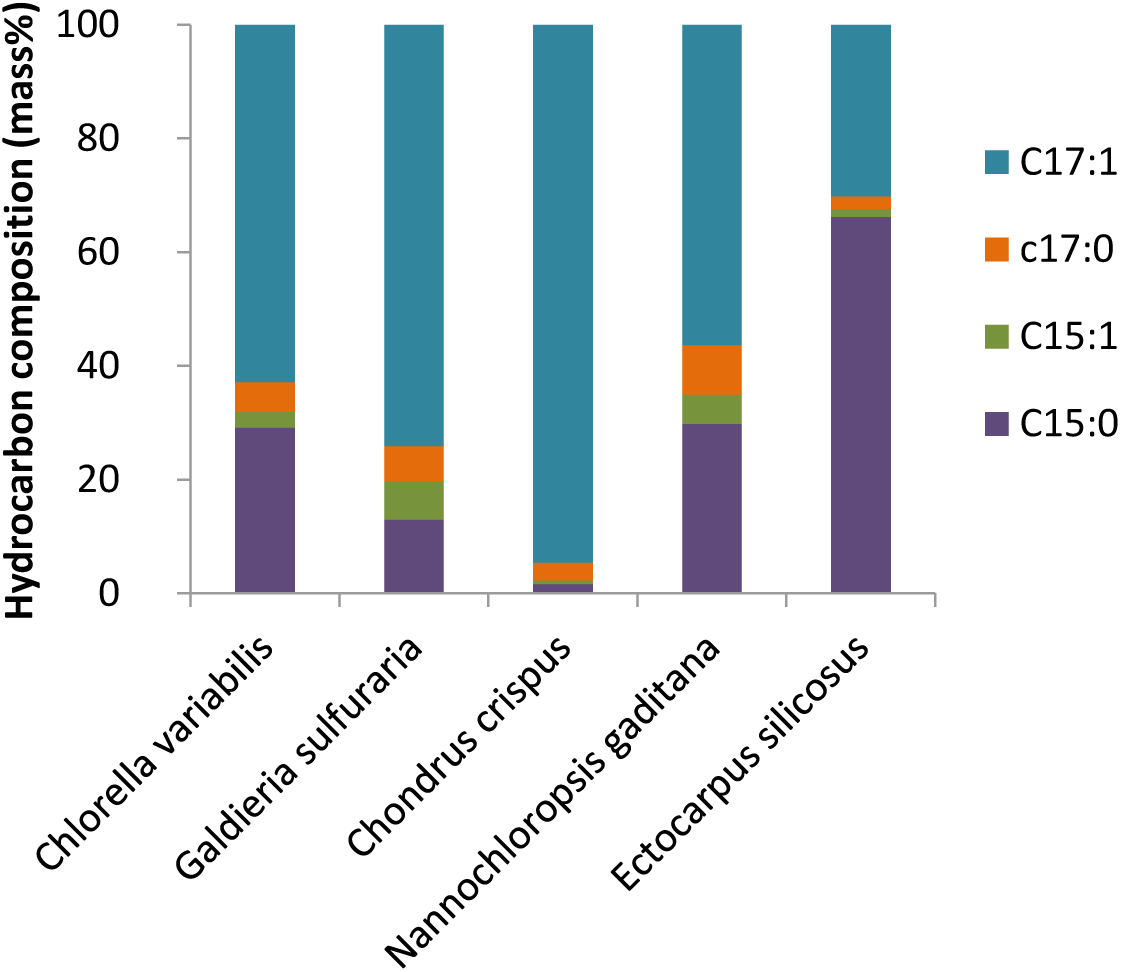
Profile of hydrocarbons produced in *E. coli* cells expressing FAP homologs. Various algal homologs of Chlorella FAP were expressed in *E. coli* and hydrocarbon composition was analysed by GC-MS after transmethylation of whole cells. Data are means of 3 independent cultures. See Figure S1 for amounts of HC formed in *E. coli* cells (mg mL^-1^).

### Identification of a new reservoir of putative FAPs

To provide a wider picture of the occurrence and evolution of putative FAP photoenzymes within algal groups and to increase the reservoir of FAPs for future biotechnological purposes, a large phylogenetic analysis of GMC oxidoreductases sequences was conducted. We used GMC oxidoreductases retrieved from public databases, from sequenced algal genomes (Blaby-Haas and Merchant, 2019) and from the Tara Ocean project (de Vargas *et al*., 2015). Tara data gave a unique opportunity to enlarge the FAP dataset with marine algal species that may not be easy to grow under laboratory conditions and whose genome has not been sequenced. Protein sequences sharing between 50 and 33% of homology with the sequence of *Chlorella variabilis* FAP were retrieved using Basic Local Alignment Search Tool (BLAST) (**Supplemental table S1 and table S2**). Over 500 GMC oxidoreductases were thus identified in the algal genomes and in the TARA dataset using annotations of the reconstructed genomes done by the Tara group. Additional GMC oxidoreductases selected from public databases were from various taxa including the three different kingdoms.

Molecular phylogeny confirmed that all the sequences of the FAP clade belong to algal species **(Fig. 3, Supplemental Fig. S2)**. Sequences from plants as well as other streptophytes (including charophytes) did not group with algal FAPs. Absence of FAP in charophytes indicated early loss of FAP function in streptophytes. No putative FAP sequences could be found in cyanobacteria although this group is highly represented in TARA data (de Vargas *et al*. 2015). Phylogeny within the FAP branch indicated that red algae (rhodophytes) sequences were the most basal. Interestingly, FAP sequences from secondary endosymbiosis-derived species appeared to be more closely related to FAPs of green algae (chlorophytes) than red algae. Overall, the new putative FAPs that could be identified in algae were present in a variety of algal groups, including stramenopiles (heterokonts), haptophytes and dinophytes. Logo sequence of FAPs compared to other GMC oxidoreductases exhibited conserved patterns **(Supplemental Fig. S3)**, including residues specific to FAP and thought to play a role in the catalysis such as C432 and R451 of CvFAP (Sorigué *et al*. 2017). Most eukaryotic algae harbored one putative FAP and no other GMC oxidoreductase, but a few algae showed no FAP and/or several non-FAP GMC oxidoreductases **(Fig. 4)**. Indeed, no putative FAP could be found in the sequenced algal genomes of the glaucocystophyte *Cyanophora paradoxa*, the Mamiellophyceae *Ostreoccocus, Micromonas* and *Bathycoccus*, the diatom *Thalassiosira pseudonana*. Conversely, only a few algal sequences could be found in other branches of the GMC oxidoreductase family. Existence in the diatom *Thalassiosira pseudonana* of a GMC oxidoreductase grouping with bacterial choline dehydrogenase was supported by one sequence from the sequenced genome (Tps-GMC) and one sequence from Tara (48230190). A Tara sequence annotated as a *Pelagomonas* protein (5166790) also turned out not to be located on the FAP clade. The cryptophyte *Guillardia theta* had 3 different GMC oxidoreductases in 3 different branches but none of them grouped with FAPs. *Ulva mutabilis* had 11 predicted GMCs, but only one was in the FAP clade. The ten other members of this multigene family of *Ulva* appeared to form a group close to plant GMC oxidoreductases. Although exceptions similar to these ones probably exist in algal diversity, the general picture appears to be that most algae have one GMC oxidoreductase, which groups with CrFAP and CvFAP in the phylogenetic tree.

**Figure 3.**
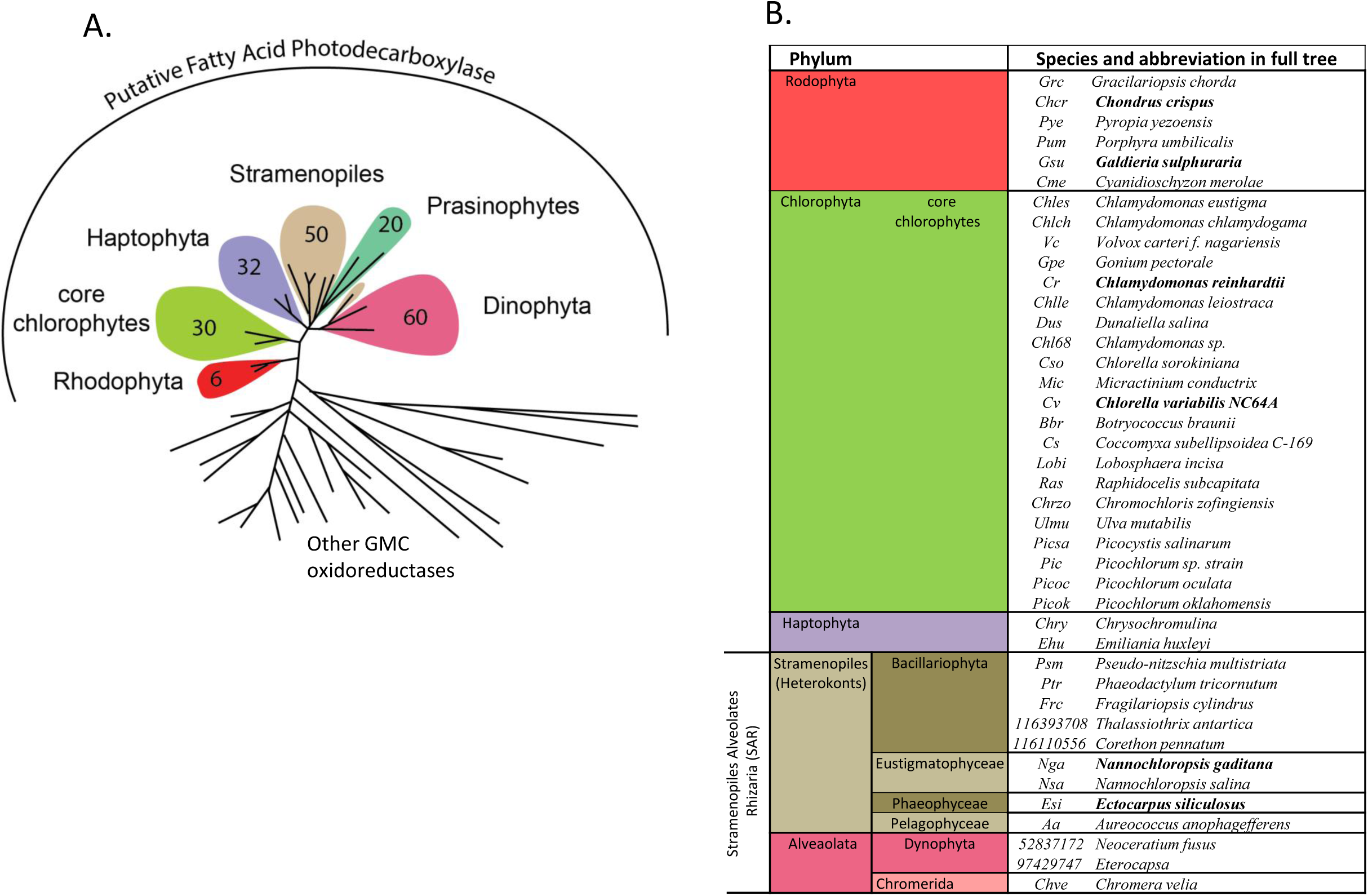
Identification of a new set of putative FAPs across algal groups. **A**, Simplified circular tree of GMC oxidoreductases showing the number of putative FAP sequences found in each group of algae. The 198 putative FAPs identified all belong to algae and have been found in TARA data (161 FAPs) and in sequenced algal genomes (37 FAPs). The tree was built using maximum likelihood algorithm using GMC oxidoreductases from various kingdoms. Annotations are focused on the branch of putative FAPs, other branches are other GMC oxidoreductases. Branches have been collapsed, full tree is available in Figure S4. **B**, Names of algae species with at least one putative FAP. For most algal groups, the number of species listed in B is lower than the one indicated in A because many species from TARA data have no annotation down to species level. When the biochemical activity of the FAP homolog is demonstrated (Sorigué *et al*., 2017 or this study), species are indicated in bold.

**Figure 4:**
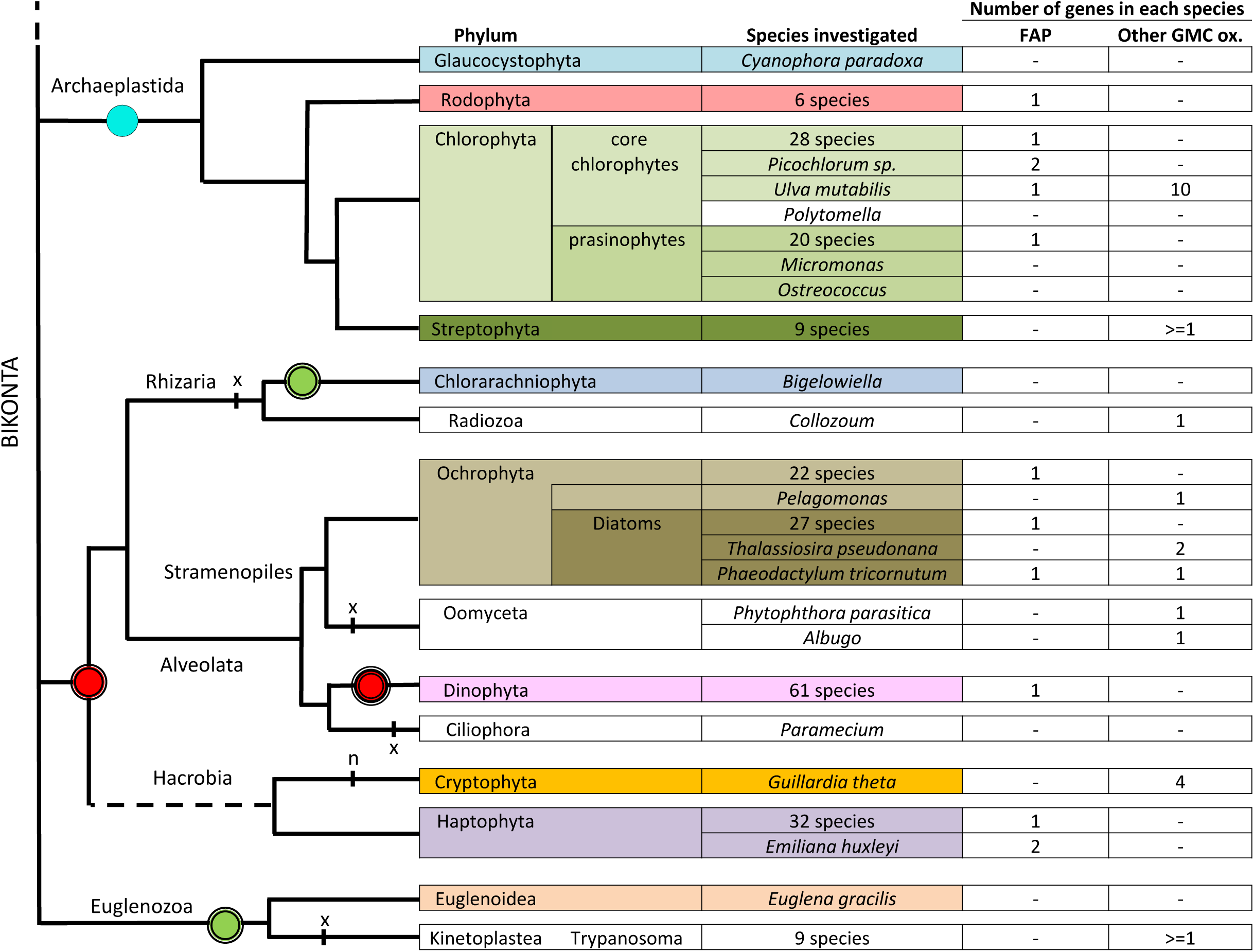
Overview of the number of FAP homologs and other GMC oxidoreductases identified in eukaryotic algae and other bikonts. In most groups, there is one FAP and no other GMC oxidoreductase. Remarkable species whose number of FAP or other GMC oxidoreductases depart from this rule are listed individually. Hyphen indicate that no protein could be identified by BLAST searches. A common indicative phylogeny is used. Photosynthetic groups or species are colored. Rounds correspond to endosymbiosis; blue round with one black circle, primary endosymbiosis; green and red rounds with two black circles, secondary endosymbiosis; red rounds with three black circles is for tertiary endosymbiosis in some (not all) Dinophyta; red or green rounds indicate red or green plastid origin respectively; n: nucleomorph; x: secondary plastid loss.

### Chlamydomonas FAP and most of its alkene products are found in the thylakoid fraction

FAP is predicted to be addressed to the chloroplast by Predalgo (Tardif *et al*., 2012), a software dedicated to the analysis of subcellular targeting sequences in green algae. This is consistent with the finding that CrFAP was found in a set of 996 proteins proposed to be chloroplastic in *Chlamydomonas* (Terashima *et al*., 2011). A broader study of putative targeting peptides using Predalgo (Tardif *et al*., 2012) and ASAFind algorithms (Gruber *et al*., 2015) indicated that FAPs from various green and red algae were largely predicted to be chloroplastic **(Supplemental Fig. S4)**. In algae with secondary plastids (i.e. containing 3 or 4 membranes), the presence of a signal peptide was consistent with a targeting to the ER or chloroplast ER (CER) membrane. Analysis performed using ASAFind, a prediction tool designed to recognize CER targeting motifs in signal peptides, indicated that such a motif was present in *Ectocarpus silicosis* and *Nannochloropsis gaditana*. Taken together, these results suggest that FAP homologs are very likely to be localized to chloroplasts in green algae, in red algae and also in at least some of the algae that acquired plastids through secondary endosymbiosis.

To consolidate these observations, subcellular fractionation of *Chlamydomonas* cells was performed **(Fig. 5A)**. Thylakoid membranes were isolated from whole cells using a sucrose gradient. Co-purification with thylakoids was followed by D1 protein (PsbA) from PSII core complex, a thylakoid membrane protein. The fact that the phosphoribulokinase (PRK) control from stroma could barely be detected in our thylakoid fraction, indicated the presence of little amount of intact chloroplasts or cells. It is thus clear that CrFAP is present in the chloroplast of *Chlamydomonas* and at least partially bound to the thylakoids. When analyzing the percentage of 7-heptadecene in total fatty acids in whole cells versus purified thylakoid membranes, a slight but significant enrichment in alkene was found in the thylakoids **(Fig. 5B)**. Using the fatty acid C16:1(3t) as a marker of the thylakoid lipids, it could be estimated that the enrichment in 7-heptadecene corresponds in fact to the localization of >90% of this compound to thylakoids **(Fig. 5C)**. These results therefore demonstrate that part of the FAP and the vast majority of the FAP product are associated to the thylakoid membranes of *Chlamydomonas*.

**Figure 5:**
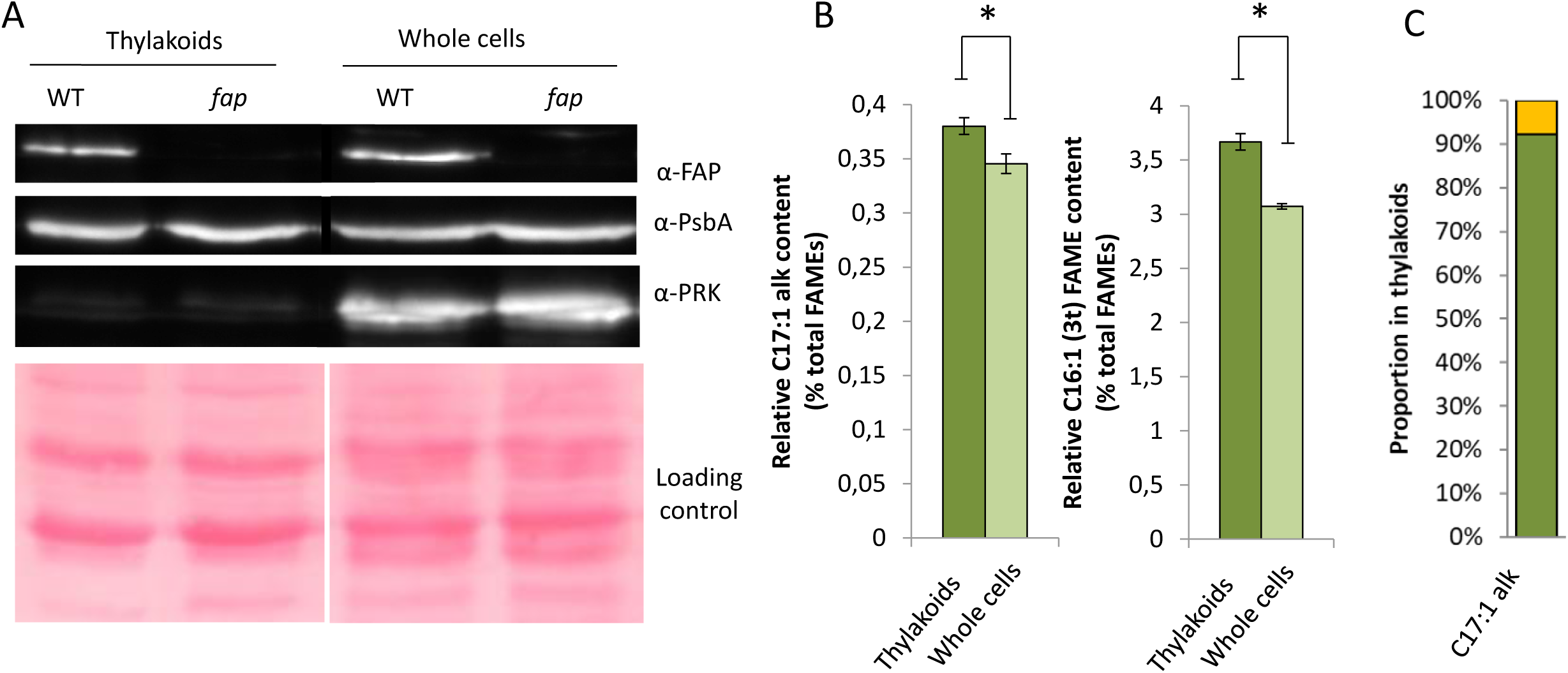
FAP and 7-heptadecene are present in the thylakoid fraction of the chloroplast. **A**, Western blot on total protein extracts from whole cells and purified thylakoid fraction. PsbA and PRK are thylakoid and stromal proteins, respectively. **B**, Relative content in 7-heptadecene and C16:1 (3t) fatty acid in whole cells and in the purified thylakoid fraction. This fatty acid almost exclusively present in thylakoids is shown for comparison. **C**, Percentage of total 7-heptadecene found in thylakoids and in the rest of the cell. Percentage was estimated using C16:1 (3t) fatty acid as a marker of thylakoids (see Material and Methods for calculation). *fap*, FAP knockout; Cp 1 to 4: complemented strains; nd: not detected. Values are mean ± SD of n=4 experiments. (*) denote p-value<0.05 in 2-sided t-test.

### 7-Heptadecene content varies with cell cycle in Chlamydomonas

The lack of FAP and HCs in chloroplasts of *C. reinhardtii* did not result in any obvious differences in the overall organization of cells or chloroplast as seen by transmission electron microscopy (TEM) **(Supplemental Fig. S5)**. To try to gather clues on FAP function, FAP transcriptomic data publicly available were mined. Transcriptomic data from Zones et al., 2015 shows that FAP has a similar expression pattern as those genes encoding proteins of the photosynthetic apparatus **(Supplemental Fig. S6)**. In order to determine whether FAP product varied with time, we monitored total fatty acids and 7-heptadecene content during a day-night cycle in synchronized *Chlamydomonas* cells. While total fatty acid content per cell increased during the day and was divided by two during cell division at the beginning of night **(Fig. 6A)**, a constant level of 7-heptadecene representing 0.45% of total fatty acids was found most of the time **(Fig. 6B)**. A significant peak (0.7%) was observed before cell division, which decreased during mitosis. This result therefore indicates that the extra-amount of HCs synthesized before cell division must be somehow lost or metabolized during cell division.

**Figure 6.**
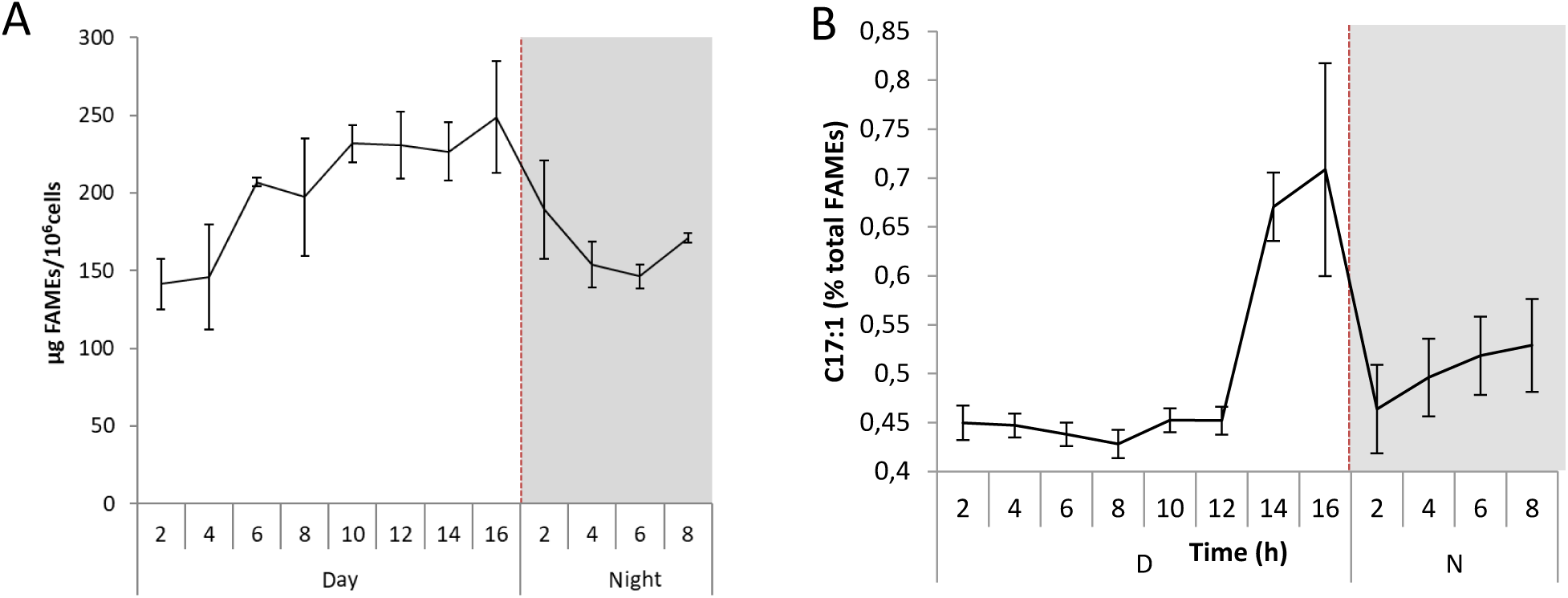
Variation of 7-heptadecene compared to total fatty acids during cell cycle. **A**, 7-Heptadecene content of cells expressed as a percent of total FAMEs. Values are mean ± SD (n=3 biological replicates). **B**, Total fatty acid content of cells during cell cycle. Amount of total fatty acids as FAMEs was analyzed by GC-MS and normalized by cell number. Data are mean ± SD of n=3 independent cultures.

### Fatty acid and membrane lipid compositions are altered in the *fap* mutant

Since 7-heptadecene content varied during cell cycle and may thus play a role in cell division, growth of the WT and *fap* strains were analyzed. Growth at 25 °C was compared in photoautotrophic conditions (mineral medium (MM)) and in mixotrophic conditions (Tris-acetate-phosphate medium (TAP)). No difference between WT and *fap* could be detected neither in growth rates nor in cell volumes under these conditions **(Supplemental Fig. S7)**. In addition, no difference between WT and *fap* strains could be observed when cells were grown under various concentrations of sodium chloride **(Supplemental Fig. S8)**. Although the lack of HCs had no effect on growth, fatty acid profile showed some differences in C16:1(9), C18:1(9), C18:3(9-12-15) **(Supplemental Fig. S9)**. Synchronized cells also showed no growth differences between WT and *fap* strains but exhibited differences in the dynamics of some fatty acid species **(Supplemental Fig. S10)**. Changes in fatty acid profiles prompted us to perform a lipidomic analysis by UPLC-MS/MS. Interestingly, it revealed that a limited set of lipid molecular species were significantly different between WT and *fap* and that were all plastidial lipids belonging to the galactolipid classes digalactosyldiacylglycerol (DGDG) and monogalactosyldiacylglycerol (MGDG) **(Fig. 7 and Supplemental Fig. S11)**. The decrease in the relative content of these galactolipid species appeared to be fully restored by complementation in the case of DGDG but not of MGDG **(Fig. 7)**. Taken together, these results show that the lack of 7-heptadecene in the *fap* mutant causes a change in thylakoid lipid composition, which is evidenced by the decrease of the relative content in at least 3 galactolipid species belonging to the DGDG class.

**Figure 7.**
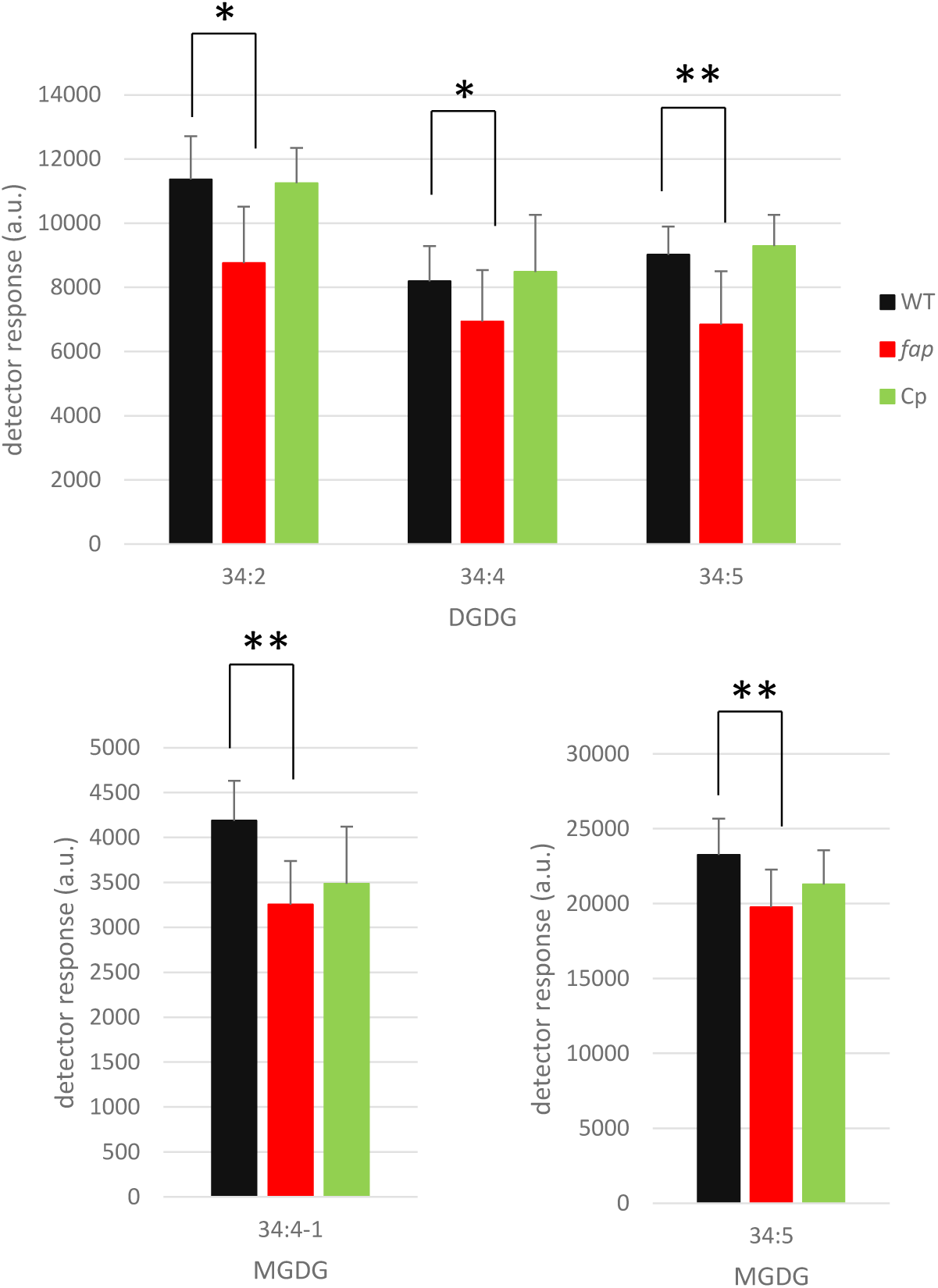
Identification of lipid molecular species significantly different between WT and *fap* strains. Relative abundance of glycerolipids was measured by LC-MS/MS analysis of total lipid extracts of whole cells. Only glycerolipid molecular species showing significant differences between WT and *fap* strains are shown here (See supplemental figure S12 for complete results). Cp, complementant strain. Lipid extracts from the three strains were loaded on a constant total fatty acid basis. Data are mean ± SD of n=9 independent cultures. MGDG34:4-1 is one of the two species of MGDG 34:1 species (which differ by 18:3 fatty acid isomers). Stars indicate significant differences according to a Mann-Whitney U-test at p<0.05 (*) or p<0.01 (**). Cells were grown in TAP medium, under 80 µmol photons m^−2^ s^−1^ in erlens.

### FAP is not strongly associated to photosynthetic complexes and lack of HCs has no effect on their organization

In cyanobacteria a role of HCs in photosynthesis has been suggested (Berla *et al*., 2015) but is controversial (Lea-Smith *et al*., 2016). In *C. reinhardtii*, there was no difference in the 77K chlorophyll fluorescence spectrum between WT, complementant and *fap* mutant, which indicated that no major changes in antenna distribution around photosystems **(Supplemental Fig. S12A)**. No difference either could be detected in photosynthesis efficiency between WT, complementant and *fap* strains grown under standard laboratory conditions **(Supplemental Fig. S12B)**. Membrane inlet mass spectrometry (MIMS) experiments conducted to quantify O_2_ exchange showed no difference in respiration and photosynthesis rates between the two genotypes **(Supplemental Fig. S12C)**. Native electrophoresis of proteins from purified thylakoids and FAP immunodetection revealed that FAP could only be detected at an apparent molecular size of the monomeric FAP **(Fig. 8)**, indicating no strong association to proteins of photosynthetic complexes. Besides, no difference in organization of photosynthetic complexes between WT and *fap* could be seen on the native protein electrophoresis.

**Figure 8.**
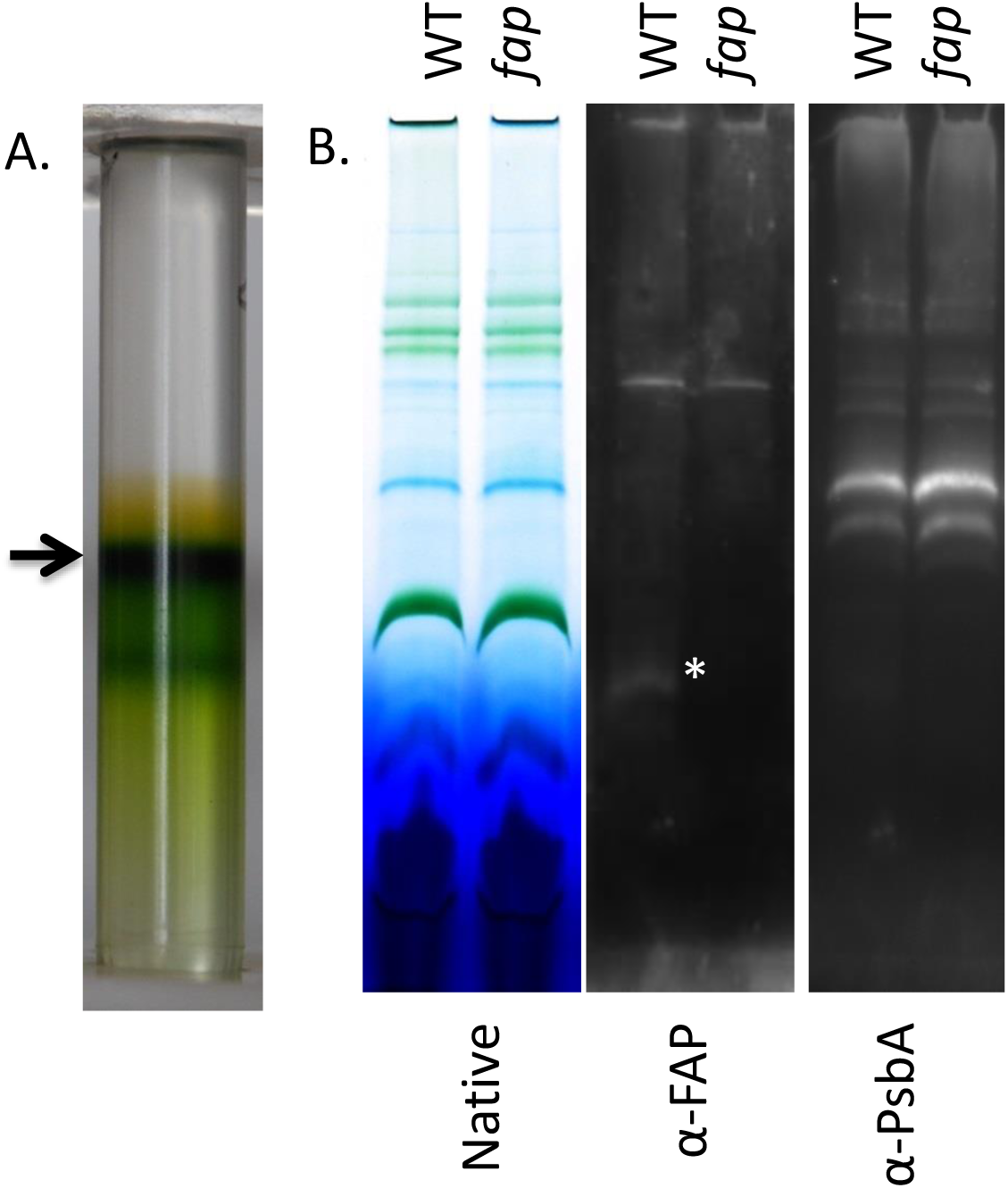
Thylakoid purification and immunoblot analysis. **A**, Thylakoid purification using sucrose density gradient. The thylakoid fraction collected is indicated by an arrow. **B**, Blue native polyacrylamide gel of solubilized proteins (0.5% digitonine, 0.5% α-DM) and corresponding immunodetection. WT: Wild type strain, *fap*: FAP knock out strain. * indicates FAP band.

### Photosynthesis is affected under light and cold stress in the *fap* mutant

Lack of HCs in the *fap* strain did not cause changes in the photosynthesis activity under standard growth conditions. However, since significant modifications in the composition of membrane lipids could be detected, we explored harsher conditions to challenge further photosynthetic membranes. We chose to investigate chilling temperatures because cold is well-known to affect both membrane physical properties and photosynthesis. Using multicultivator in turbidostat mode, we first stabilized cultures at 25 °C under medium light (200 µmol photons m^−2^ s^−1^), electron transfer rate (ETR) showed no difference **(Fig. 9A)**. When cooling down the culture to 15 °C and after 3 days of acclimation, both ETR and 77 K chlorophyll fluorescence spectra still showed no differences **(Fig. 9B**,**C) (Fig. 9C)**. After one day at a lower light intensity (50 µmol photons m^−2^s^−1^), the maximal PSII yield was equal for all the strains but ETR was lower for the mutant when measured at high light intensities **(Fig. 9D)**. Interestingly, longer acclimation to this condition (3 days) led to the disappearance of this phenotype.

**Figure 9.**
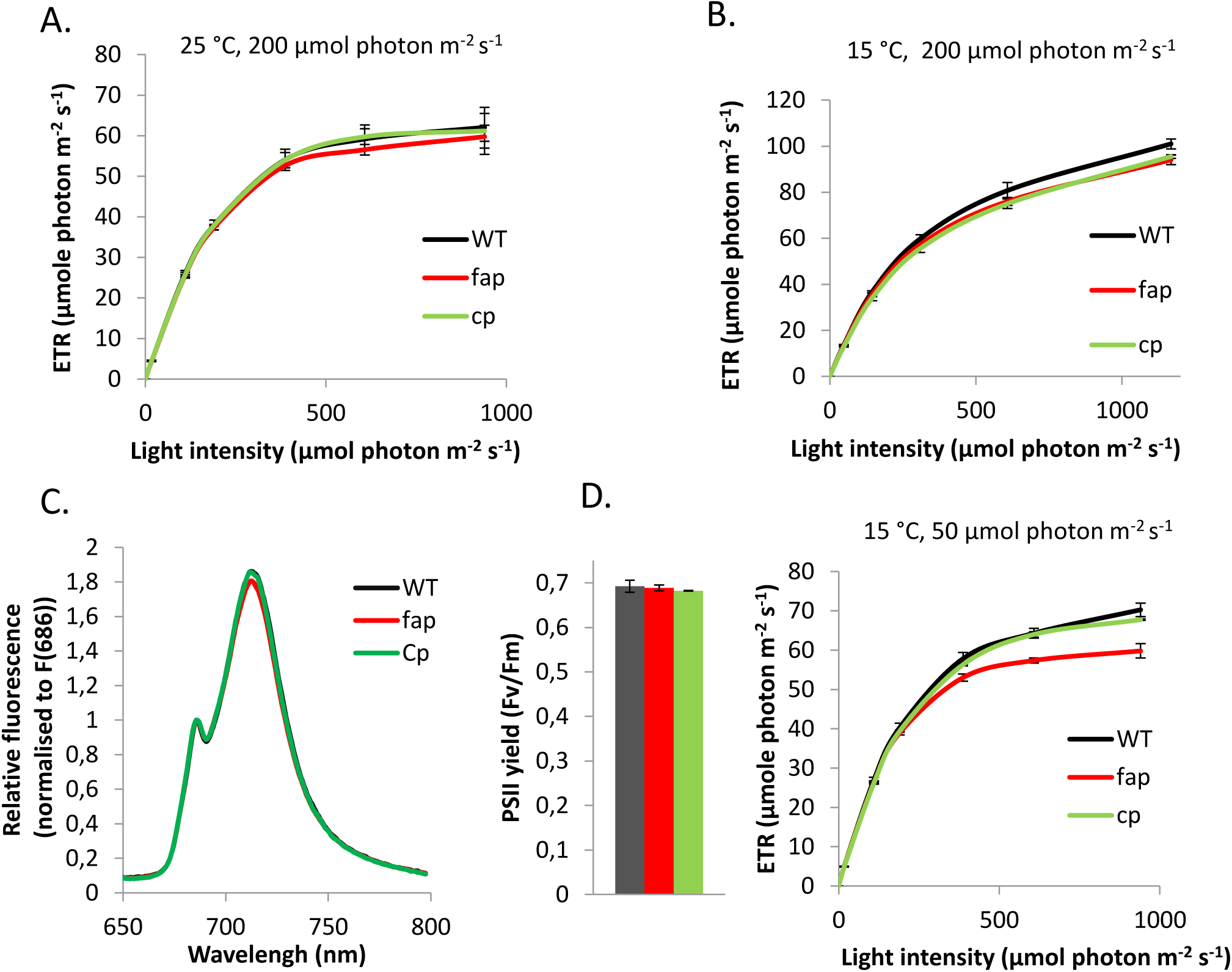
Photosynthetic acclimation to cold conditions in WT and *fap* strains. Electron transfer rate (ETR) at various light intensities for cells grown in photoautotrophic conditions at 25 °C (**A**) and 15 °C (**B**). **C**, 77K fluorescence spectrum for cold conditions (15 °C). **D**, ETR and PSII yield at 15 °C and lower light than in A,B. Data are mean ± SD of n=3 independent cultures.

In order to provide support for a possible link between HCs and cold acclimation, 7-heptadecene content was quantified under various growth temperatures. Relative HC content in cells clearly increased under cold conditions **(Fig. 10)**. As expected, an increase in the relative content in polyunsaturated species occurred upon cold treatment **(Supplemental Fig. S13)**, but no difference in the dynamics of fatty acid remodeling was observed between WT and *fap* strains.

**Figure 10.**
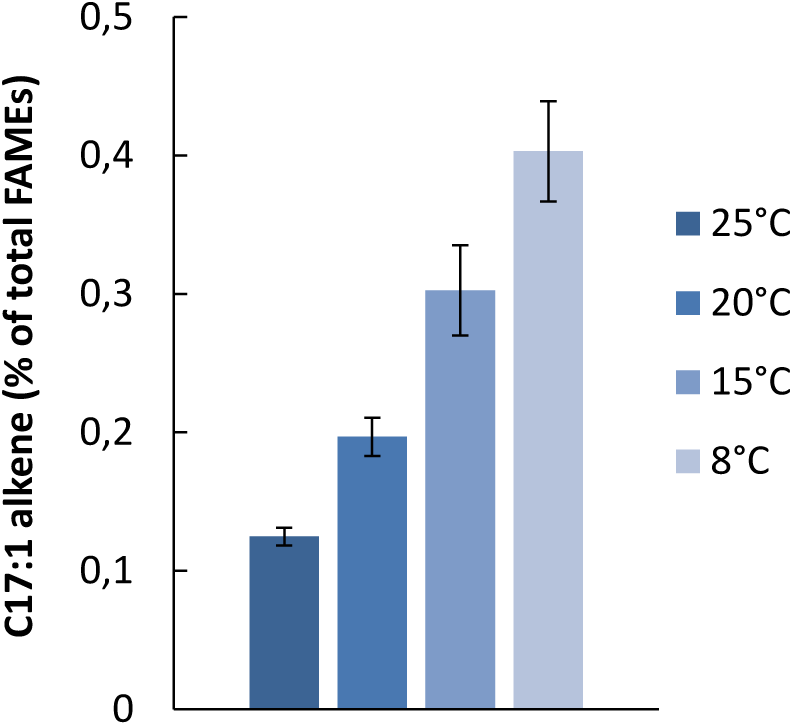
Hydrocarbon amount in cells cultivated at various temperatures. Hydrocarbon and fatty acid content was analyzed by GC-MS after transmethylation. Cells were grown in photobioreactors, in turbidostat mode, in TAP medium under 50 µmol photons m^-2^ s^-1^. Data are mean ± SD of n=3 independent cultures.

## DISCUSSION

Here, we report the isolation and characterization of an insertional *Chlamydomonas* mutant deficient in *FAP* and we perform a phylogenetic and functional analyses of algal homologs. We show FAP and the vast majority of its 7-heptadecene product are associated to thylakoid membranes. It is also shown that the *FAP* gene is present in most algal lineages and encodes a functional fatty acid photodecarboxylase in some species of red algae, of secondary algae as well as in some macroalgae. By studying a FAP knock-out *Chlamydomonas* mutant, we provide evidence that lack of hydrocarbons is correlated with small changes in galactolipid composition but has no impact on photosynthesis and growth in *Chlamydomonas* under standard culture conditions. However, in the absence of hydrocarbons generated by FAP, the photosynthetic activity is transitorily affected during cold acclimation. The possible significance of these results for algal physiology as well as FAP function and evolution are discussed below.

### FAP and formation of HCs in algal cells

Based on the characterization of a *fap* mutant, we first show that FAP is responsible for the synthesis of all fatty acid-derived HCs found in *Chlamydomonas* cells **(Fig. 1)**. Our result clearly demonstrates that the fatty acid photodecarboxylase activity measured *in vitro* for CrFAP (Sorigué *et al*., 2017) is not a promiscuous secondary activity and indeed corresponds to a genuine biological activity, namely the light-driven synthesis of 7-heptadecene from *cis*-vaccenic acid **(Fig. 11)**. Also, the *fap* knockout line shows that no other enzyme is able to synthesize 7-heptadecene in *Chlamydomonas*. Besides, the fact that HC production was found to be correlated with the quantity of FAP present in complemented lines, indicated that FAP is a limiting factor for 7-heptadecene production *in vivo*. Thus, the putative lipase activity that must be acting upstream of FAP to generate the free *cis-*vaccenic acid is not limiting in the pathway.

**Figure 11.**
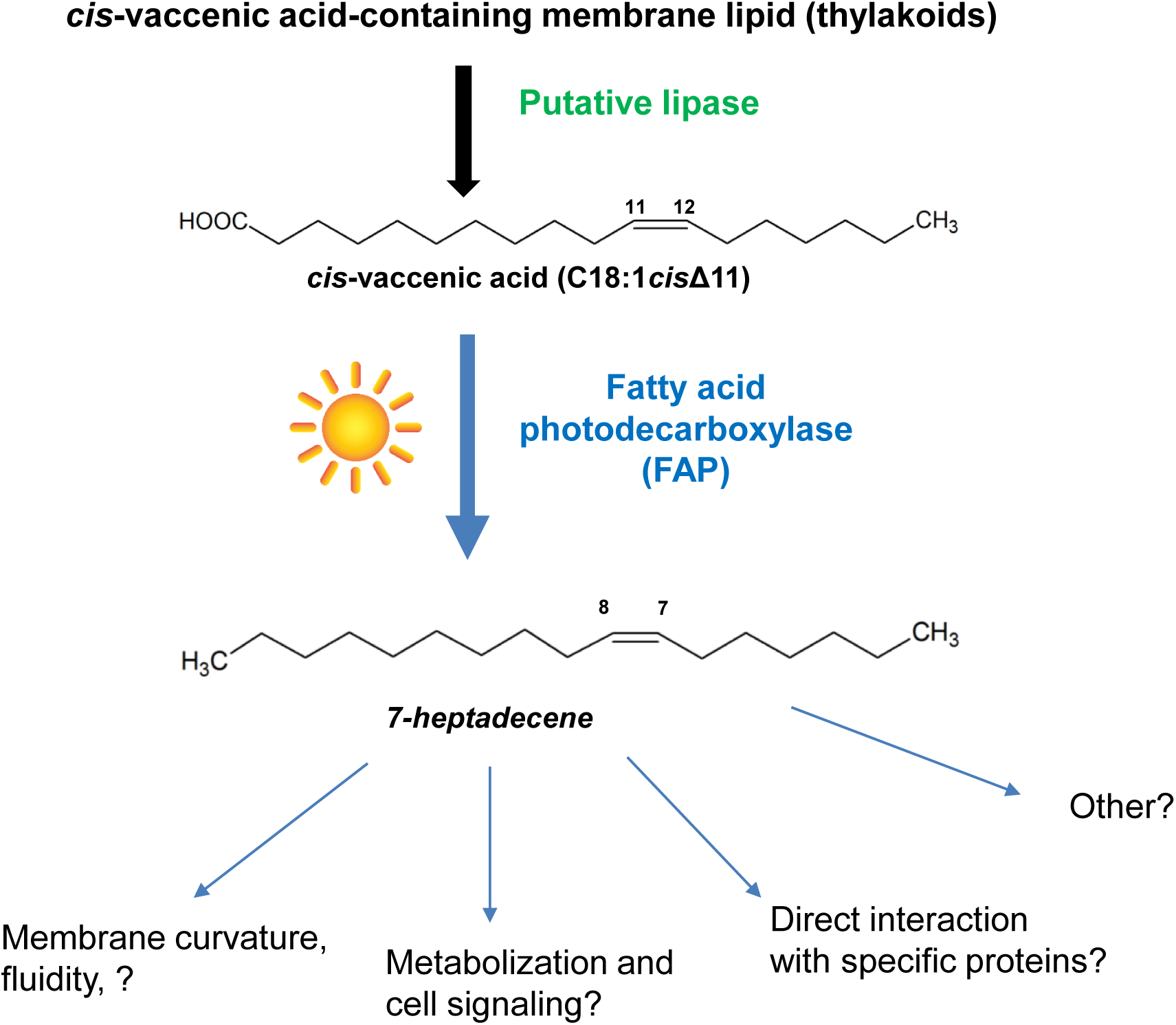
Proposed pathway for hydrocarbon formation from fatty acids in *Chlamydomonas* and putative roles. The only fatty acid-derived hydrocarbon in *C. reinhardtii* (7-heptadecene) is generated from *cis*-vaccenic acid by FAP only when cells are exposed to light. The fatty acid precursor must be released from a thylakoid lipid by an unknown lipase. The 7-heptadecene FAP product may play several roles in thylakoid membranes depending on temperature and light conditions.

### Localization of FAP and role in membranes

Based on subcellular fractionation and anti-FAP antibodies, we show here that FAP is able to associate to thylakoid membranes **(Fig. 5)**. This result is consistent with the predicted plastid localization and with the fact that thylakoid membranes harbored 90% of the 7-heptadecene product. Presence of plastid transit peptides in FAPs seems to be a general rule in green and red algae (primary plastids) and is also predicted for some secondary endosymbiosis-derived algae **(Supplemental Fig. S4)**.

Considering that HCs are hydrophobic compounds, it is not surprising that in *Chlamydomonas* the 7-heptadecene is mostly located where it is produced. In a work on cyanobacterial mutants devoid of fatty acid-derived HCs, it has been suggested that HCs are located in membranes and may play a role in cell division (Lea-Smith *et al*., 2016). In the proposed cyanobacterial model, integration of HCs into the lipid bilayer would be responsible of membrane flexibility and curvature. HCs may play a similar role in thylakoid membranes of green algae. The fact that the percentage of 7-heptadecene in the total fatty acids stays rather stable during a 16-hour day, indicates that HC production follows lipid production during cell growth, except just before mitosis **(Fig. 6B)**. In addition, the ratio of HCs to FAMEs decreased at the beginning of night, when cells are dividing, indicating that some HCs are lost during cell division. A simple mechanism which could explain HC loss during cell division involves enrichment in HCs at breaking points of plastidial membranes before cell division, exclusion from these membranes during division and loss to the gas phase of the culture due to HC volatility **(Supplemental Fig. 6B)**.

HCs might thus impact local flexibility of algal plastidial membranes and participate in lipid membrane remodeling during cell division. However, under standard culture conditions, the presence of HCs is apparently not critical for chloroplast structure **(Supplemental Fig. 5)**, cell size and cell division rate **(Supplemental Fig. 7)**. A role of cyanobacterial HCs in resistance to salt stress has also been suggested (Yamamori *et al*., 2018). In *Chlamydomonas*, contrary to what has been shown in cyanobacteria, no difference could be detected in growth under increasing salt concentrations **(Supplemental Fig. 8)**. One could thus hypothesize that even if HCs are produced in chloroplast and accumulated in thylakoids, their function might be different from that in cyanobacteria. It is also possible that laboratory culture conditions used for *Chlamydomonas* (this study) are far from natural growth conditions where HCs may be necessary. Alternatively, compensation mechanism for HC loss may operate differently in *Chlamydomonas* and in cyanobacteria. In *Chlamydomonas*, part of this mechanism may involve changes in membrane lipid composition. Interestingly, lipidomic analysis under standard growth conditions unraveled specific changes in DGDG molecular species **(Fig. 7)** but no other significant differences in other class of lipids **(Supplemental Fig. S11)**. Taken together, these results suggest that in *Chlamydomonas* HCs play no crucial role in cell division and growth under standard conditions. Cells may adapt to a lack of HCs by some changes in the composition of membranes, which could specifically involve some DGDG galactolipids. Alternatively, or in addition to this proposed effect on properties of the membrane lipid phase, it cannot be ruled out that 7-heptadecene may act locally to disrupt or enhance some specific protein-protein interactions, or may play a yet to be defined role, such as acting as a signaling molecule or its precursor **(Fig. 11)**.

### FAP and photosynthetic membranes

The fact that the FAP gene expression that follows that of photosynthesis genes in day-night cycles, the likely localization of FAP in plastids of green and red algae as well as in some secondary algae, and the localization of part of FAP and almost all its alkene product in *Chlamydomonas* thylakoids point toward a role of FAPs in the photosynthetic function of algal cells. This idea is strongly reinforced by the conservation of the FAP-encoding gene in many eukaryotic algae but not in non-photosynthetic protists **(Fig. 3 and Fig. 4)** and in *Polytomella*, an algae that has kept some of its plastidial function but lost photosynthesis (Smith and Lee 2014). As standard culture conditions did not allow to reveal any photosynthesis phenotype in *Chlamydomonas fap* mutant **(Supplemental Fig. S12)**, more challenging conditions involving colder temperatures and variations in light intensity were tested. These experiments have revealed a difference between WT and *fap* mutant in the photosynthesis activity measured under high light during acclimation to cold **(Fig. 9)**. Interestingly, colder temperatures are correlated with increased HCs **(Fig. 10)** while fatty acid profiles follow the same trend in WT and *fap* strains **(Fig. S13)**. Taken together, these observations indicate that adaptations in membrane lipid composition compensate partly for the loss of HCs in standard growth conditions but not in harsher conditions such as cold temperatures.

### Conservation of FAP in algae

According to molecular phylogeny **(Fig. 3, Fig. 4)**, FAP proteins appear to be specific to algae and highly conserved in many algae species. A noticeable exception is the Mamiellophyceae class of the green algae. Algae is a common denomination that gathers photosynthetic eukaryotes which mainly live in aquatic environments. This polyphyletic group includes organisms derived from a first endosymbiosis as well as organisms derived from a secondary or even tertiary endosymbiosis. However, a functional FAP can be found in chlorophytes (green algae), rhodophytes (*Chondrus* and *Galdieria*) and stramenopiles (in the phaeophyceae *Ectocarpus* and the Eustigmatophyceae *Nannochloropsis*) as proven by heterologous expression in *E. coli* of the corresponding identified FAPs **(Fig. 2)**. FAP activity was therefore conserved during secondary endosymbiotic event(s) that gave rise to the red lineage. Moreover, FAP activity is not specific to the unicellular state as FAPs were also functional in the pluricellular algae (macroalgae) *Ectocarpus silicosus* and *Chondrus crispus*. Considering homology of sequences, FAP function is thus expected to be present in most algal phyla, including haptophytes and dinophytes (dinoflagellates). Importantly, some aminoacid residues that are likely to be involved in fatty acid substrate stabilization or photocatalysis, such as CvFAP Arg451 or Cys432 (Sorigué *et al*., 2017) are strictly conserved in the 198 putative FAPs **(Supplemental Fig. S3)**. This observation reinforces the idea that all the putative FAPs identified in this work have the ability to photo-produce HCs from fatty acids.

FAP neofunctionalization from GMC oxidoreductases may have occurred early during evolution of algae, almost concomitantly with the very first endosymbiosis shared by green and red algae. No GMC could be found in glaucophytes, which may indicate that this event has occurred after separation of glaucophytes from red and green algae. However it should be noted that so far only one complete glaucophyte genome is available. Absence of FAP in charophytes indicates early loss of FAP function in streptophytes. Phylogeny points out that FAPs from secondary endosymbiosis lineages are more closely related to core chlorophytes than rhodophytes. FAP could thus be one of the genes that was inherited from green algae by horizontal gene transfer (Moustafa *et al*., 2009).

Concerning the conservation of FAP activity, it should be noted that the FAPs selected for heterologous expression produced various HCs profiles **(Fig. 2)**. For example, *Chondrus* FAP showed high specificity for C18:1 fatty acid producing 95% of C17:1 alkene, while *Ectocarpus* FAP produced 70% of C15:0 alkane. This indicates that the algal biodiversity contains FAPs which are more selective or more active on shorter chain fatty acids than FAPs of *Chlorella* and *Chlamydomonas*. FAPs with different properties may be useful for biotechnological application aiming to enhance the production of short chain volatile HCs by microbial cell factories (Moulin *et al*., 2019).

In conclusion, the results presented here show that FAP activity is conserved beyond green microalgae and identify a big reservoir of FAPs that may be useful for biotechnological applications. It also provide some important clues for future studies aiming at unravelling the exact role of the FAP photoenzyme in eukaryotic algae.

## MATERIALS AND METHODS

### Strains and culture conditions

The *fap* mutant and its corresponding wild-type strain of *C. reinhardtii* were ordered from the CLiP library (Li *et al*. 2016). Upon reception, strains were plated on Tris-acetate-phosphate (TAP) medium and streaked to allow formation of single colonies. For each strain, after 1-week growth in the dark, three single-clone derived colonies were randomly chosen for characterization. Wild type strains are CC-4533 cw15 mt^-^ for mating type minus and CC-5155 cw15 mt^+^ (Jonikas CMJ030 F5 backcross strain) [isolate E8] for mating type plus. Mutant LMJ.RY0402.226794 was used in this study, which is predicted to harbor a first insertional cassette in coding sequence of Cre12.g514200 encoding FAP. A second insertion in the line LMJ.RY0402.226794 was predicted in Cre14.g628702. To remove this side mutation we backcrossed the mutant strain to CC-5155. Analysis of one full tetrad showed 2 progeny strains resistant to paromomycin which were mutated in *FAP* gene. The region of Cre14.g628702 was amplified by PCR and sequenced for the 4 progeny strains of the tetrad. No insertion was actually found, therefore a potential insertion at this locus was ruled out and the initial prediction by the CLip project was not accurate and this could happen due to the strain mixing during handling of large number of clones (Li *et al*. 2016). Work on mutant strains was conducted on one parental isolated strain with the mutation from LMJ.RY0402.226794 and the 2 mutants of the full tetrad from the backcross with CC-5155. This 3 strains are thereafter named *fap-1, fap-2, fap-3* respectively. Wild type (WT) strains were parental strain CC5155 (WT-1) and single colony-derived lines of background strain CC4533 (WT-2, WT-3).For liquid culture experiments, cells were grown in 24 deep well plates of 25 mL under 100 µmol photons m^-2^ s^-1^ with constant shaking at 25 °C. Cells were grown in TAP or minimal medium (MM) (Hoober 1989) for mixotrophic and autotrophic conditions, respectively. Cell growth was followed using a cell counter Multisizer (Beckman Coulter). For day night cycle experiment, cells were cultivated autotrophically in 1L-photobiorectors in turbidostat mode (Dang *et al*. 2014; Sorigué *et al*. 2016) (OD_880nm_ at 0.4) under 16 hours of light (40 µmol photons m^-2^ s^-1^) 8 hours of dark at 25 °C. For photosynthesis analysis, cells were grown autotrophically in 80 mL photobioreactors (multicultivator, Photon Systems Instruments) with turbidostat module (OD_680nm_ at 0.8). Conditions were 25 °C, medium light (200 µmol photons m^-2^ s^-1^), or 15 °C medium light or 15 °C low light (50 µmol photons m^-2^ s^-1^). All cultures were done under ambient air.

### Complementation of the *fap* mutant

Construct for complementation of knockout strain for *FAP* gene was carried out using pSL-Hyg vector containing an *AphVII* cassette conferring hygromycin resistance **(Supplemental Fig. S14)**. This vector allowing nuclear transformation was kindly provided by Pr. Steven Ball (University of Lille, France). WT copy of the *FAP* gene was obtained by PCR of WT genomic DNA using primers Cr-F and Cr-R (This and all other primer sequences were shown in **Supplemental Table S3**). It was cloned into TOPO-XL vector. pSL-Hyg vector and *FAP* gene were digested with *Eco*RV and *Spe*I and ligated. Then, the vector was linearized with *Pvu*I and was electroporated into the *fap* strains. Level of complementation was verified by immunoblot to assess quantity of protein and by transmethylation of whole cells to assess quantity of HCs.

### SDS PAGE and Immunodetection

Cells (10–15 mL) were harvested by centrifugation at 3,000 *g* for 2 min. Pellets were then frozen in liquid nitrogen and stored at -80 °C until use. Pellets were resuspended in 400 mL 1% (w) SDS and then 1.6 mL acetone precooled to -20 °C was added. After overnight incubation at -20 °C, samples were centrifuged (14,000 rpm, 10 min, 4 °C). Supernatant was removed and used for chlorophyll quantification using SAFAS UVmc spectrophotometer (SAFAS). Pellets were resuspended to 1 mg chlorophyll mL^-1^ in LDS in the presence of NuPAGE reducing agent (ThermoFischer) and loaded on 10% (w/v) PAGE Bis-Tris SDS gel. To load equal protein amounts for immunoblot analysis, protein contents were estimated by Coomassie Brilliant Blue staining of the gel using an Odyssey IR Imager (LICOR). After gel electrophoresis, proteins were transferred to nitrocellulose membranes for 75 min at 10 V using a semi-dry set up. Membranes were blocked in TBST, milk 5% (w/v) overnight at 4 °C then incubated at room temperature in the presence of the following antibodies: anti-Cyt f, anti-AtpB, anti-PsaD, anti-PsbA, anti-LHCSR3 (Agrisera), or anti-FAP (see below). After 2 hour incubation, primary antibody was removed by rinsing 3 times in TBST, and a peroxidase-coupled secondary antibody was added for at least 1 h. Luminescence was detected with a Gbox imaging system (Syngene).

### Production of anti-CrFAP antibodies

Codon-optimized synthetic gene encoding *C. reinhardtii FAP* (Sorigué *et al*. 2017) was cloned into the pLIC7 expression vector, allowing the production of a recombinant FAP fused to TEV-cleavable His-tagged *Escherichia coli* thioredoxin. Production was performed in the *E. coli* BL21 Star (DE3) strain initially grown at 37 °C in TB medium. Induction was initiated at an OD_600nm_ of 0.8 by adding 0.5 mM isopropyl b-D-thiogalactoside (IPTG), cultures were then grown at 20 °C. Following overnight incubation, cells were centrifuged and protein was purified as described previously (Sorigué *et al*. 2017). Purity of the purified protein was controlled on SDS-PAGE and it was brought to a final concentration of 2 mg mL^-1^ using an Amicon-Ultra device (Millipore). Polyclonal antibodies against FAP were raised in rabbits (ProteoGenix, Schiltigheim, France).

### Analysis of hydrocarbons and fatty acids

For quantification of HCs, about one hundred million cells were pelleted by centrifugation in glass tubes. Transmethylation was conducted by adding 2 mL of methanol containing 5% (v/v) sulfuric acid to the cell pellet. Internal standards (10 µg of hexadecane and 20 µg of triheptadecanoylglycerol) were added for quantification. Reaction was carried out for 90 min at 85 °C in sealed glass tubes. After cooling down, one mL of 0.9% (w/v) NaCl and 500 µL of hexane were added to the samples to allow phase separation and recovery of fatty acid methyl esters (FAMEs) and HCs in the hexane phase. Samples were mixed and then centrifuged to allow phase separation. Two µL of the hexane phase was injected in the GC-MS/FID. Analyses were carried out on an Agilent 7890A gas chromatographer coupled to an Agilent 5975C mass spectrometer (simple quadrupole). A Zebron 7HG-G007-11 (Phenomenex) polar capillary column (length 30 m, internal diameter 0.25 mm, and film thickness 0.25 mm) was used. Hydrogen carrier gas was at 1 mL min^-1^. Oven temperature was programmed with an initial 2-min hold time at 35 °C, a first ramp from 35 to 150 °C at 15 °C min^-1^, followed by a 1-min hold time at 170 °C then a second ramp from 170 to 240 °C at 5 °C min^-1^ and a final 2-min hold time at 240 °C. The MS was run in full scan over 40 to 350 amu (electron impact ionization at 70 eV), and peaks of FAMEs and HCs were quantified based on the FID signal using the internal standards C17:0 FAME and hexadecane, respectively.

### Chlorophyll fluorescence measurements and MIMS analysis

Chlorophyll fluorescence measurements were performed using a pulse amplitude-modulated fluorimeter (Dual-PAM 100) upon 15-min dark-adaptation under continuous stirring. Detection pulses (10 mmol photons m^-2^ s^-1^ blue light) were supplied at a 100-Hz frequency. Basal fluorescence (F_0_) was measured in the dark prior to the first saturating flash. Red saturating flashes (6,000 mmol photons m^-2^ s^-1^, 600 ms) were delivered to measure F_m_ (in the dark) and F_m’_ (in the light). Electron transfer rate (ETR) was measured with a saturating flash after 2 to 3 minutes of illumination at a given light intensity. PSII maximum yields were calculated as (F_m_ – F_0_)/ F_m_ and PSII yield for each light intensity was calculated from (F_m’_ – F)/F_m’_. ETR was calculated as the product of light intensity and PSII yield. MIMS was used to measure gas exchange as described previously (Burlacot *et al*. 2018).

### Transmission electron microscopy

Cells were grown photoautotrophically in photobioreactors under 40 µmol photons m^-2^ s^-1^ in turbidostat (OD_880nm_ at 0.4). The algal cells were collected by centrifugation (1 500 *g*, 1 min) and were immediately fixed with 2.5% (v/v) glutaraldehyde in 0.1 M, pH 7.4 sodium cacodylate buffer for two days at 4 °C. They were then washed by resuspending 5 min three times in the same buffer. Samples were post-osmicated with 1% (w/v) osmium tetroxyde in cacodylate buffer for 1 h, dehydrated through a graded ethanol series, and finally embedded in monomeric resin Epon 812. All chemicals used for histological preparation were purchased from Electron Microscopy Sciences (Hatfield, USA). Ultra-thin sections for transmission electron microscope (90 nm) were obtained by an ultramicrotome UCT (Leica Microsystems GmbH, Wetzlar, Germany) and mounted on copper grids and examined in a Tecnai G2 Biotwin Electron Microscope (ThermoFisher Scientific FEI, Eindhoven, the Netherlands) using an accelerating voltage of 100 kV and equipped with a CCD camera Megaview III (Olympus Soft imaging Solutions GmbH, Münster, Germany).

### Isolation of thylakoids and native PAGE

Thylakoids were isolated according to the protocol described previously (Chua *et al*. 1975). All steps were performed on ice or at 4 °C with as little light as possible. Briefly, cells were pellet and resuspended in 8 mL 25 mM, 5 mM MgCl_2_, 0.3 M sucrose, with a protease inhibitor cocktail for plant cell and tissue extracts (Sigma P 9599). Cell were disrupted with French press at a pressure of 6000 Psi. They were collected by centrifugation (1000*g*, 10 min) and washed first in 5 mM HEPES, 10 mM EDTA, 0.3 M sucrose and then in 5 mM HEPES, 10 mM EDTA, 1.8 M sucrose. Sucrose gradient was 0.5 M sucrose (5 mL), 1.3 M sucrose (2 mL) and 1.8 M sucrose initially containing thylakoids (5 mL). After ultracentrifugation (274 000*g*, 1h), thylakoids were collected between 0.5 and 1.3 M sucrose. They were washed with 5 mM HEPES, 10 mM EDTA and resuspended at 1 mg mL^-1^ chlorophyll for subsequent SDS-PAGE analysis. For non-denaturing conditions, thylakoids were resuspended in NativePAGE sample buffer (Life technologies) at 1 mg mL^-1^ chlorophyll, thylakoids were solubilized for 30 min on ice in the same volume of 1% (w/v) n-Dodecyl-alpha-D-Maltoside, 1% (w/v) digitonine (0.5 mg mL^-1^ chlorophyll and 0.5% (w/v) n-Dodecyl-alpha-D-Maltoside, 0.5% (w/v) digitonine final). For each sample, 20 µL were then loaded with 2 µL of G-250 sample additive (Life technologies) on 4-16% (w/v) NativePAGE gels (Life technologies). Cathode running buffer (Life technologies) was supplemented with 0.02% (w/v) G-250 for two-thirds of the migration, and with 0.002% (w/v) G-250 for the remaining third. Annotation of observed bands was done according to a previous publication (Pagliano *et al*. 2012). For immunoblot analysis, native gel was incubated in Tris Glycine SDS buffer, 10% (v/v) ethanol for 15 min and transferred on PVDF membrane using XCell II Blot module (25v, 1h). Immunodetection was done as described above.

Based on C16:1(3t) FAME we determine a factor of enrichment expected for a compound that would be exclusively within thylakoids (ratio EF=C16:1 (3t) FAME_whole cells_ /C16:1 (3t) FAME_thylakoids_). Considering the amount of C17:1 alkene found in thylakoids, we calculated the expected content in whole cells which equals C17:1 alk_thylakoids_*ratio EF. This calculated value for thylakoids was divided by the value for whole cells determined experimentally, which gives the proportion of C17:1 alkane that is present in thylakoids.

### Lipidomic analysis by UPLC-MS/MS

Lipid molecular species analysis was done by Ultra Performance Liquid Chromatography coupled with tandem Mass Spectrometry (UPLC-MS/MS). Lipids were first extracted with a modified hot isopropanol method. Briefly, *C. reinhardtii* cells were harvested by centrifugation at 4000 rpm, 2 min in glass tubes. Pellet were immediately resuspended in 1 mL of hot isopropanol (85 °C) containing 0.01% (w/v) butylated hydroxytoluene (BHT). Sealed tubes were heated at 85 °C for 10 minutes to inactivate lipases. Internal standards were added. Lipids were then extracted in 3 mL methyl tert-butyl ether(MTBE) with a phase separation with 1 mL of water. Organic phase was collected and aqueous phase was washed with an additional mL of MTBE. Organic phases were evaporated under a gentle nitrogen stream and resuspended in 500 μL of a mixture of acetonitrile/isopropanol/ammonium acetate 10 mM (65:30:5, v/v/v). Lipid molecular species were analyzed on an ultimate RS 3000 UPLC system (ThermoFisher, Waltham, MA, USA) connected to a quadrupole-time-of-flight (QTOF) 5600 mass spectrometer(AB Sci ex, Framingham, MA, USA) equipped with a duo-spray ion source operating in positive mode. Lipid extracts were first separated on a Kinetex™ (Kinetex, Atlanta, GA, USA) C182.1×150 mm 1.7 μm column (Phenomenex, Torrance, CA, USA). Two solvent mixtures, acetonitrile-water (60:40, v/v) and isopropanol-acetonitrile (90:10, v/v), both containing 10 mM ammonium formate at pH 3.8, were used as eluent A and B respectively. The elution was performed with a gradient of 32 min; eluent B was increased from 27 to 97% in 20 min then maintained for 5 min, solvent B was decreased to 27% and then maintained for another 7 min for column re-equilibration. The flow-rate was 0.3 mL min^-1^ and the column oven temperature was maintained at 45 °C. Lipid identification was based on retention time and on mass accuracy peaks from the MS survey scan compared with theoretical masses and fragment ions from MS/MS scan. Relative quantification was achieved with multiquant software (AB Sciex) on the basis of intensity values of extracted masses of the different lipids previously identified. Detector response was normalized by the quantity of FAME previously measured by GC-MS for each sample.

### Pigment quantification

Cell pellets were re-suspended 1 mL of methanol. After at least 1 hour at -20 °C, debris were pelleted by centrifuging and supernatants were analyzed using a spectrophotometer measuring absorbance at 450, 653, 666 and 750 nm. Concentration in different pigments are calculated according to the following formula (Ritchie *et al*., 2008; Strickland 1968) using the dilution factor (DF):

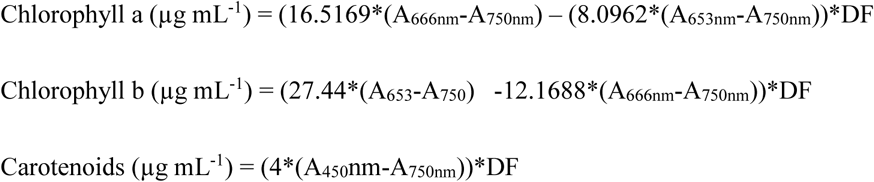

### Phylogenetic analysis and logo sequences

The CvFAP protein sequence was used as bait and blasted against different databases using tBLASTn or BLASTp (including NCBI, Phytozome, Fernbase, and Unigene TARA ocean databases). Sequences from the BLAST were pooled with reference sequences from a previous tree of the GMC oxidoreductase superfamily (Zamocky *et al*. 2004). Alignment of sequences was done with Muscle algorithm (Edgar 2004) and viewed with Seaview software. Selection of conserved sites was done enlarging Gblock with apparent conserved sites. A set of 226 conserved positions were used for tree construction using maximum of likelihood algorithm (PhyML, with LG algorithm) with 100 replicates for bootstrap analysis. Annotation of the tree was down using annotation data provided by TARA. FAP Logo sequence was based on 35 sequences including at least one sequence of each taxa from the phylogeny. GMC logo sequence was based on sequences of non-FAP GMC oxidoreductases (Zamocky *et al*. 2004). Alignment of sequences was done using using Muscle algorithm and viewed with Seaview software. Construction of Logo sequences was done using WebLogo (https://weblogo.berkeley.edu/logo.cgi).

### Heterologous expression of FAP in *E*.*coli*

The FAP homologs from *Galdieria sufuraria, Chondrus crispus* and *Ectocarpus silicosus* were codon-optimized and synthetized then cloned in pLIC07 as described before for CvFAP (Sorigué *et al*. 2017). *FAP* gene from *Nannochloropsis gaditana* was directly amplified from cDNAs. Briefly, total RNAs were extracted and the *NgFAP* was amplified by PCR from cDNA obtained by reverse transcription using primers Ng-F and Ng-R described in Supplemental Table S3 Gene prediction from NCBI was wrong as sequencing proved the absence of a predicted intron. There is thus a STOP codon in the middle of the predicted protein and actual protein sequenced stop at DEERKGGWFNGLLGRKQKAAT. Potential transit peptides were removed for better expression in *E. coli* and N-terminal sequences were as follows: GFDRSREFDYVIVGGG for *Galdieria*; SSEAATTYDYIIVGGG for *Chondrus*; LQSVSMKAPAAVASSTYDYIIVGGG *for Nannochloropsis*; SMSVAEEGHKFIIIGGG for *Ectocarpus. E. coli* were cultivated in Terrific broth medium at 37 °C until OD_600nm_ reached 0.8. Expression was then induced with 0.5 mM IPTG and transferred at 22 °C under 100 µmol photons m^-2^ s^-1^.

### 77K fluorescence emission spectra

Low temperature spectra where measured on whole cells at 77K using a SAFAS Xenius optical fiber fluorescence spectrophotometer (Dang *et al*, 2014). 200 µL of light-adapted cell suspension, was frozen in a liquid nitrogen bath cryostat. The excitation wavelength used was 440 nm and detection wavelength ranged from 600 to 800nm with a 5nm split. Fluorescence emission spectra where all normalized to the 686 nm signal.

## ACKNOWLEDGEMENTS

We thank Dr. Olivier Vallon for help with analysis of some sequenced algal genomes and useful discussions. Thanks are due to Dr. Quentin Carradec and Dr. Patrick Wincker for helping with access to TARA sequences. Help of Dr. Philippe Ortet and Emmanuelle Billon with analysis of TARA sequences is also acknowledged.

## FIGURE LEGENDS

**Figure S1.**
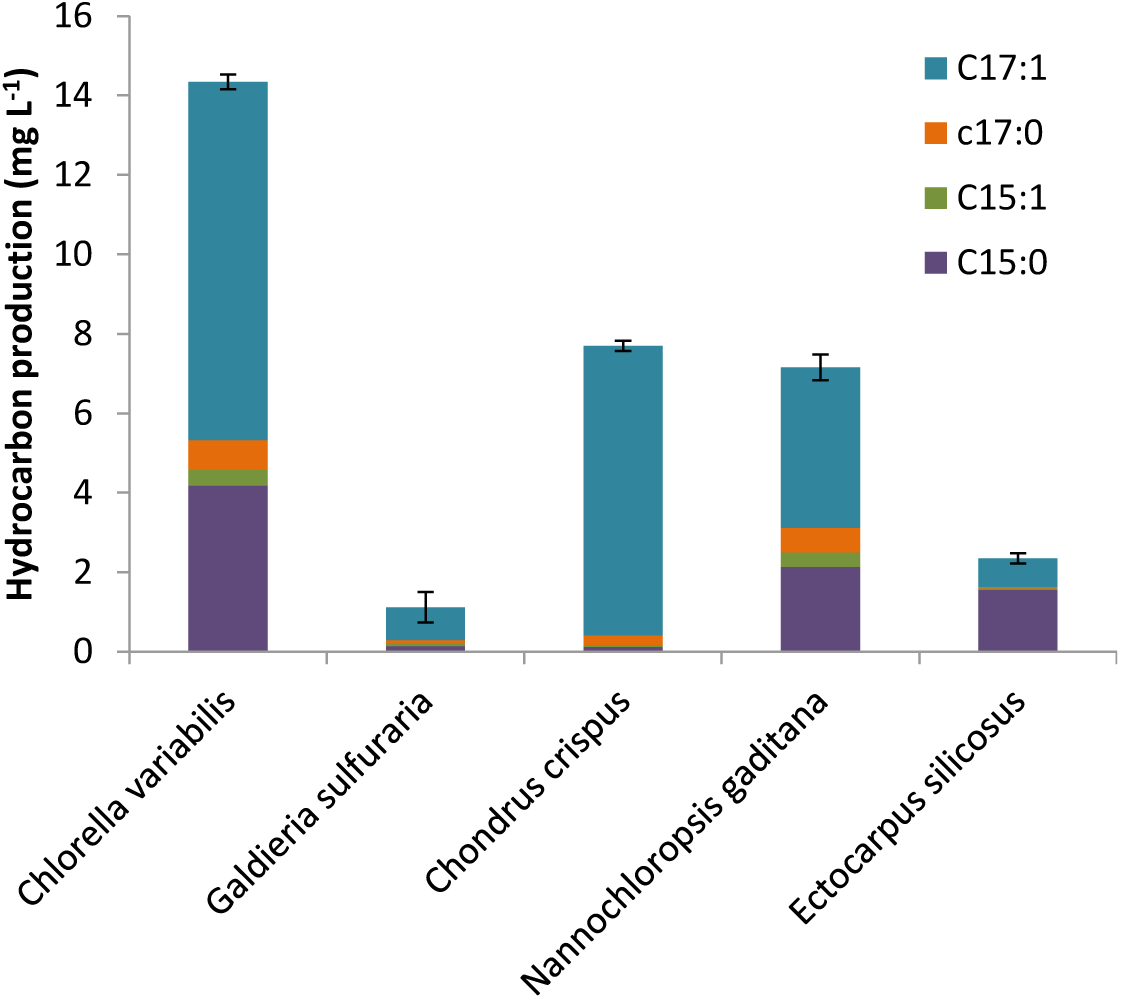
Amount of hydrocarbons produced in *E. coli* strains expressing FAPs of various origins. Hydrocarbons produced in *E. coli* cells were extracted with organic solvents and analyzed by GC-MS (means ± SD; n=3 biological replicates).

SupplementalFig. 2: phylogenetictree(7 printedpages)

**Figure S2.** Phylogeny of GMC oxidoreductase superfamily. Tree of GMC sequences built using maximum of likelihood algorithm. Each branch is defined by the lowest taxonomic indication gathering all the sequences present in the branch. When biochemical activity is demonstrated capital letters indicate it : AOX, alcohol oxidase; FAP, fatty acid photodecarboxylase; CBQ, cellobiose dehydrogenase; CHD, choline dehydrogenase; COX, cholesterol oxidase; GlucDH, glucose dehydrogenase; GOX, glucose oxidase, HNL hydroxymandelonitrile lyase. When indicated in brackets, function has been shown by phylogeny approach but is not yet supported by activity assay.

**Figure S3.**
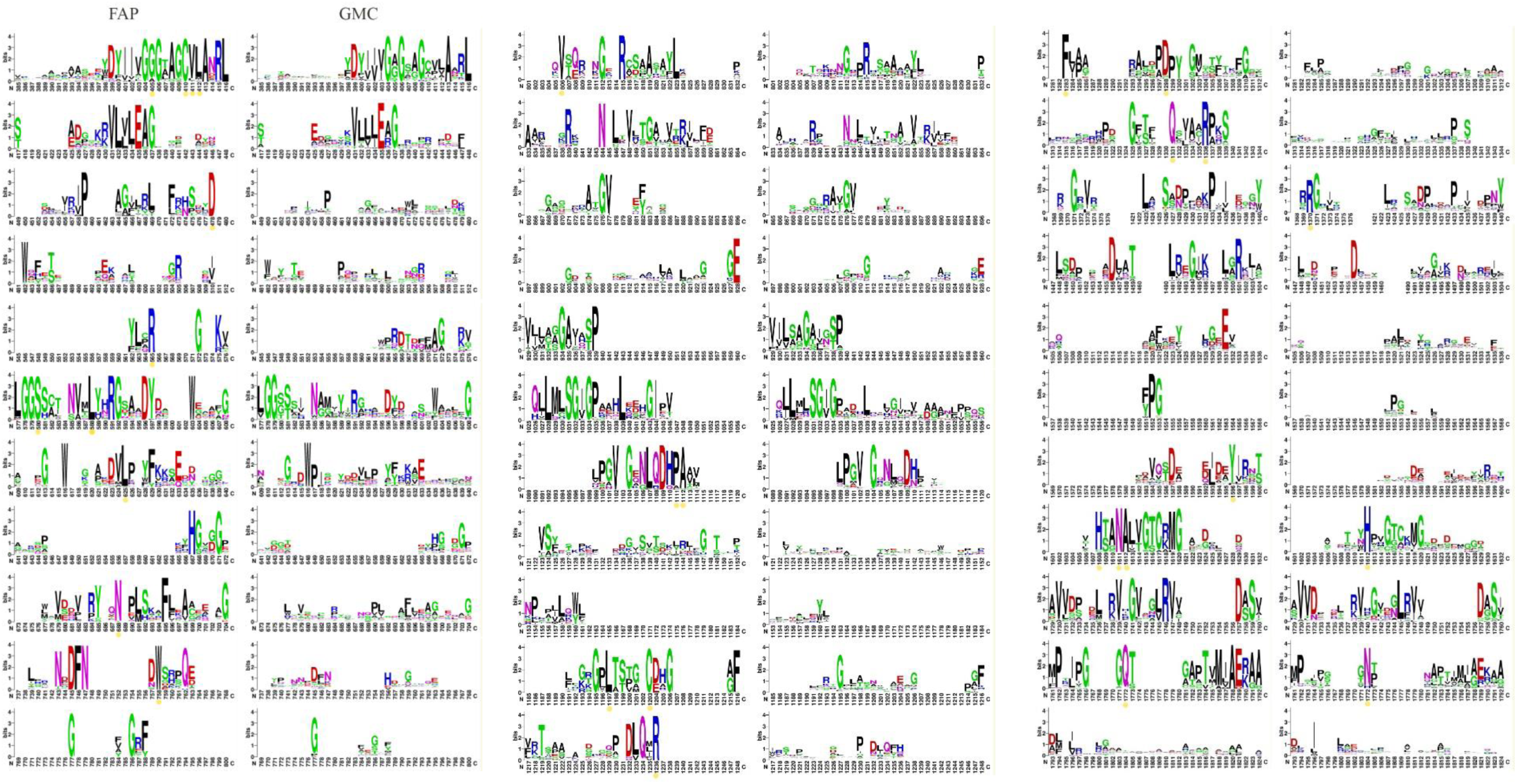
Logo sequences for FAPs and other GMC oxidoreductases. Sequences used for alignments are in supplemental table S1. Yellow stars indicate conserved residues specific to FAPs. Residues C432 and R451 of *Cv*FAP active site are indicated by a yellow star underlined in red.

**Figure S4.**
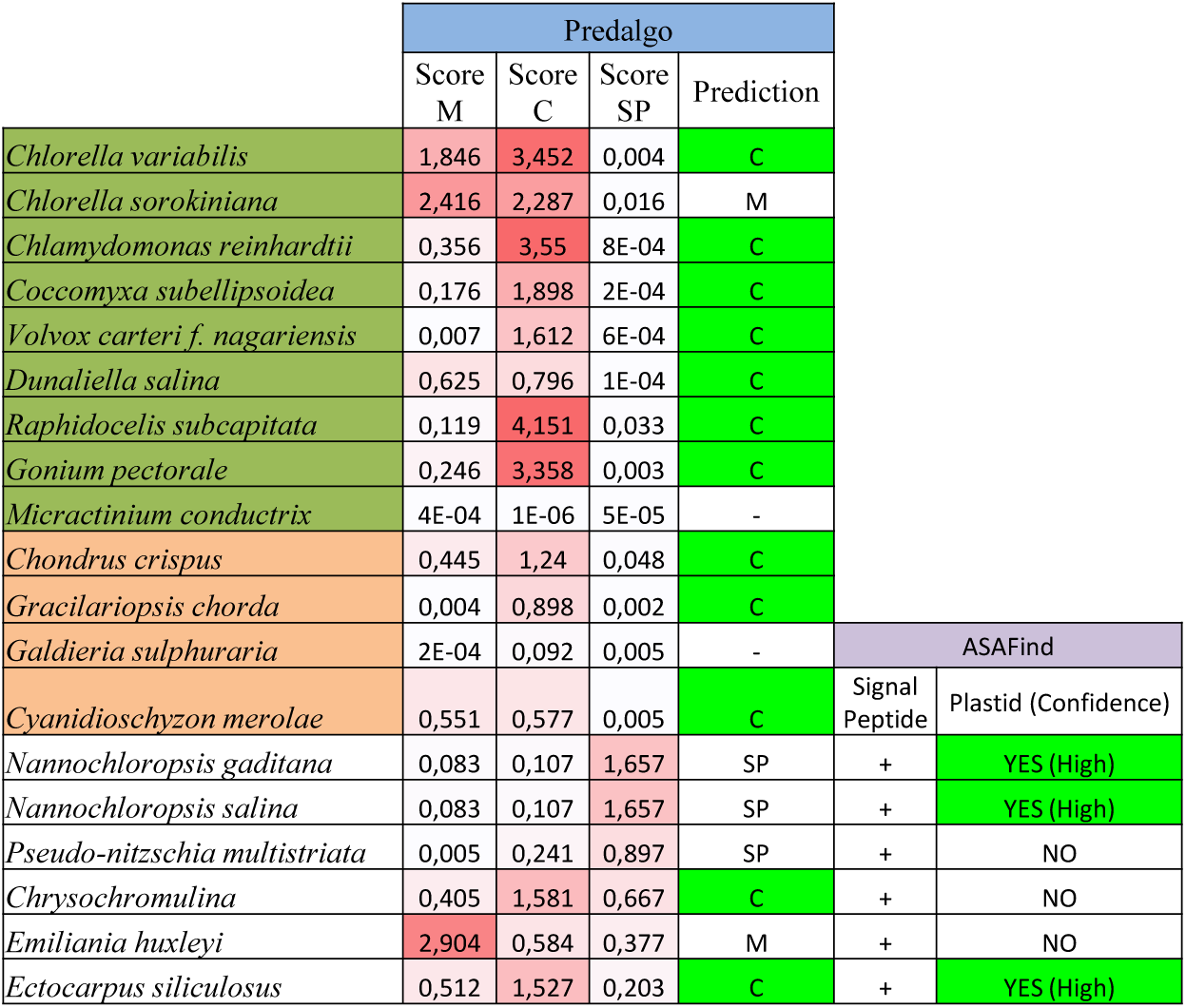
FAP is predicted to be localized in plastids in many algae. Sequences from various algae with sequenced genomes were analyzed with two algorithms adapted to algal sequences. ASAFind has been developed specifically for algae with secondary plastids. Scores are indicated and colored in red when significant. M: mitochondria, C: chloroplast, SP: secretory pathway. For secondary endosymbiosis algae, ASAFind results are presented with presence of transit peptide according to SignalP and confidence of plastidial localization.

**Figure S5.**
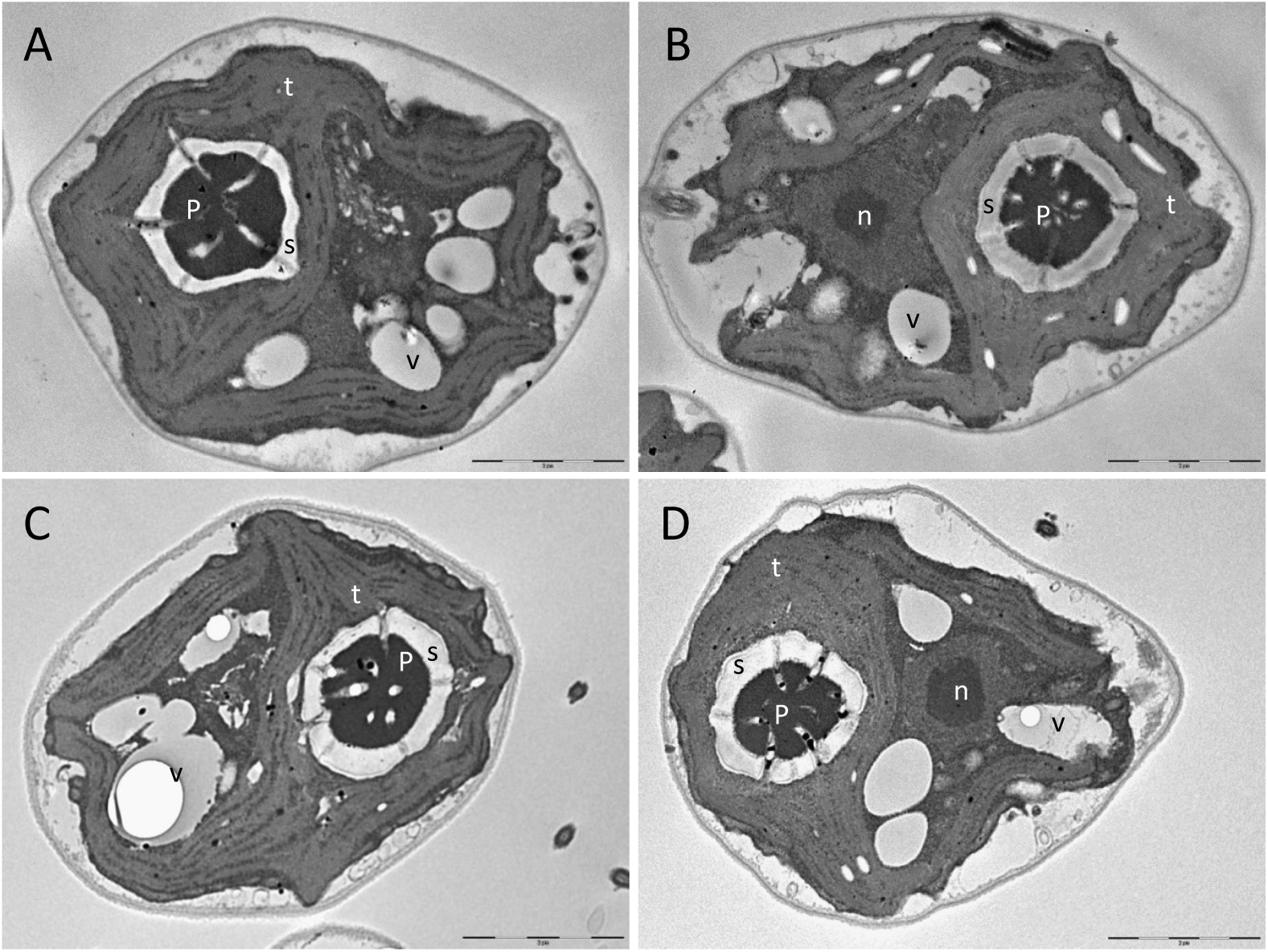
Ultrastructure of C. reinhardtii wild type and *fap* strains. Transmission electron microscopy of wild type (A,B) and fap strain (C,D). Thylakoids (t), nucleus (n), vacuoles (v), pyrenoïd (p) and starch (s). Scale bar: 2 µm.

**Figure S6.**
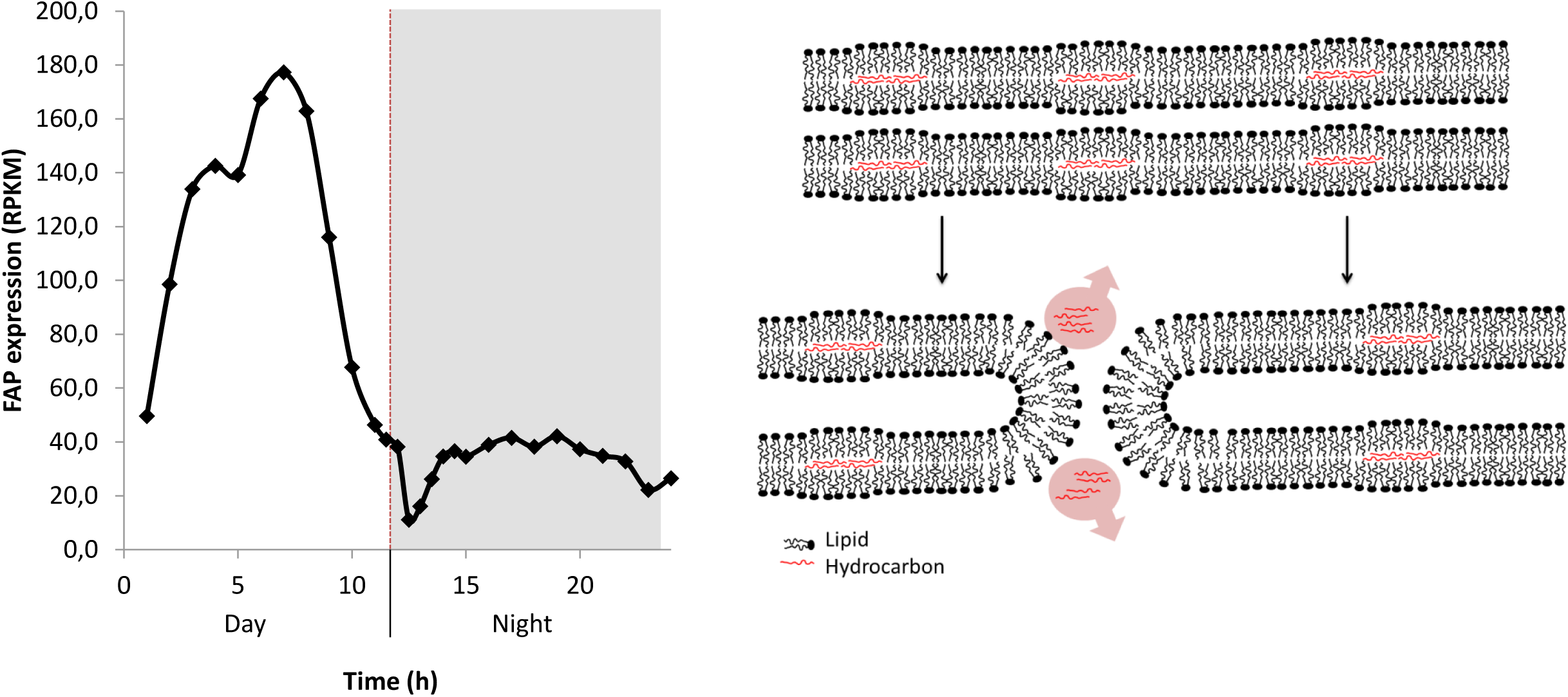
FAP gene expression during day-night cycle and hypothetical mechanism that may explain HC loss. **A**, FAP expression from transcriptomic data (from Zones et al., 2015). RPKM: reads per kilobase of transcript per million reads mapped. **B**, Proposed model for HC loss in *Chlamydomonas* chloroplasts during cell division. Alternatively, HCs could be metabolized.

**Figure S7.**
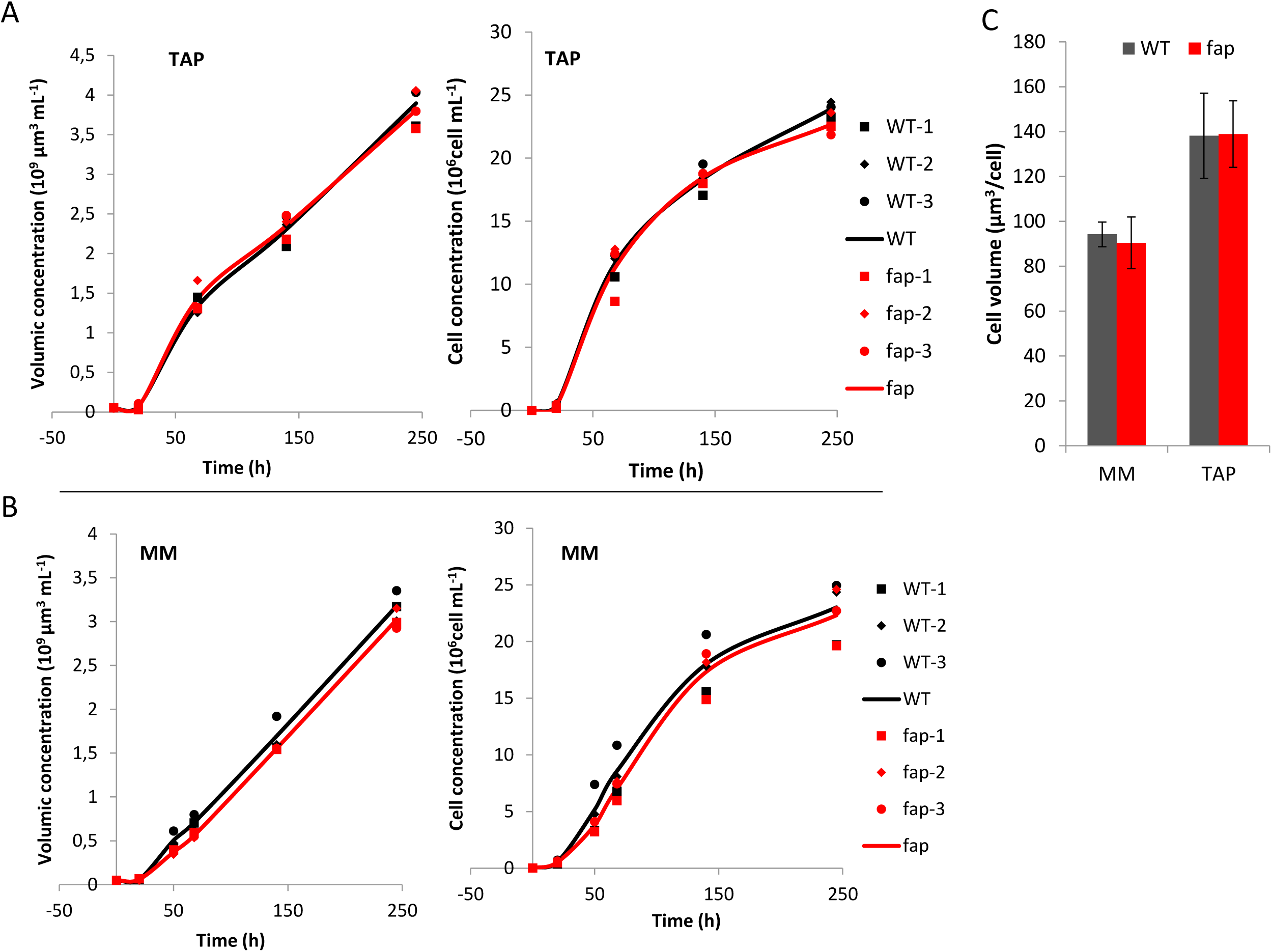
Growth curves and cell volume of wild type and *fap* strains. A, Growth on acetate medium (TAP). B, Growth on minimal medium (MM). Data are expressed in cell volume (left) or cell number (right). Data points shown are from 3 independent cultures. Curves show the average trend. C, Cell volume of wild type and fap strains after 6 days of growth in minimal medium (MM) or acetate medium (TAP). Values are means ± SD (n = 3 independent cultures).

**Figure S8:**
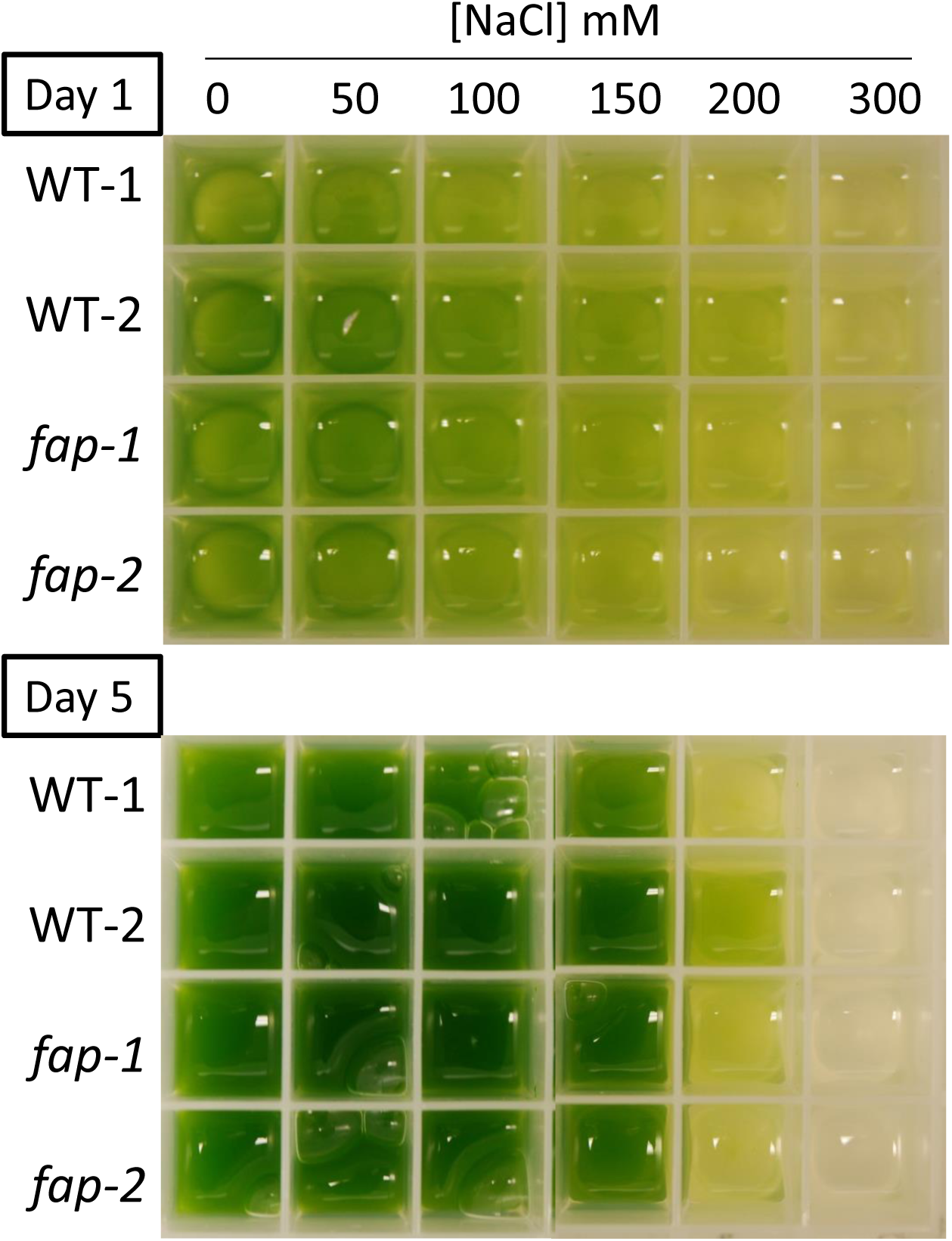
Growth of wild type and *fap* strain using various concentrations of salt. Cultures in acetate medium (TAP) were exposed 1 or 5 days to salt concentrations from 0 to 300 mM NaCl.

**Figure S9.**
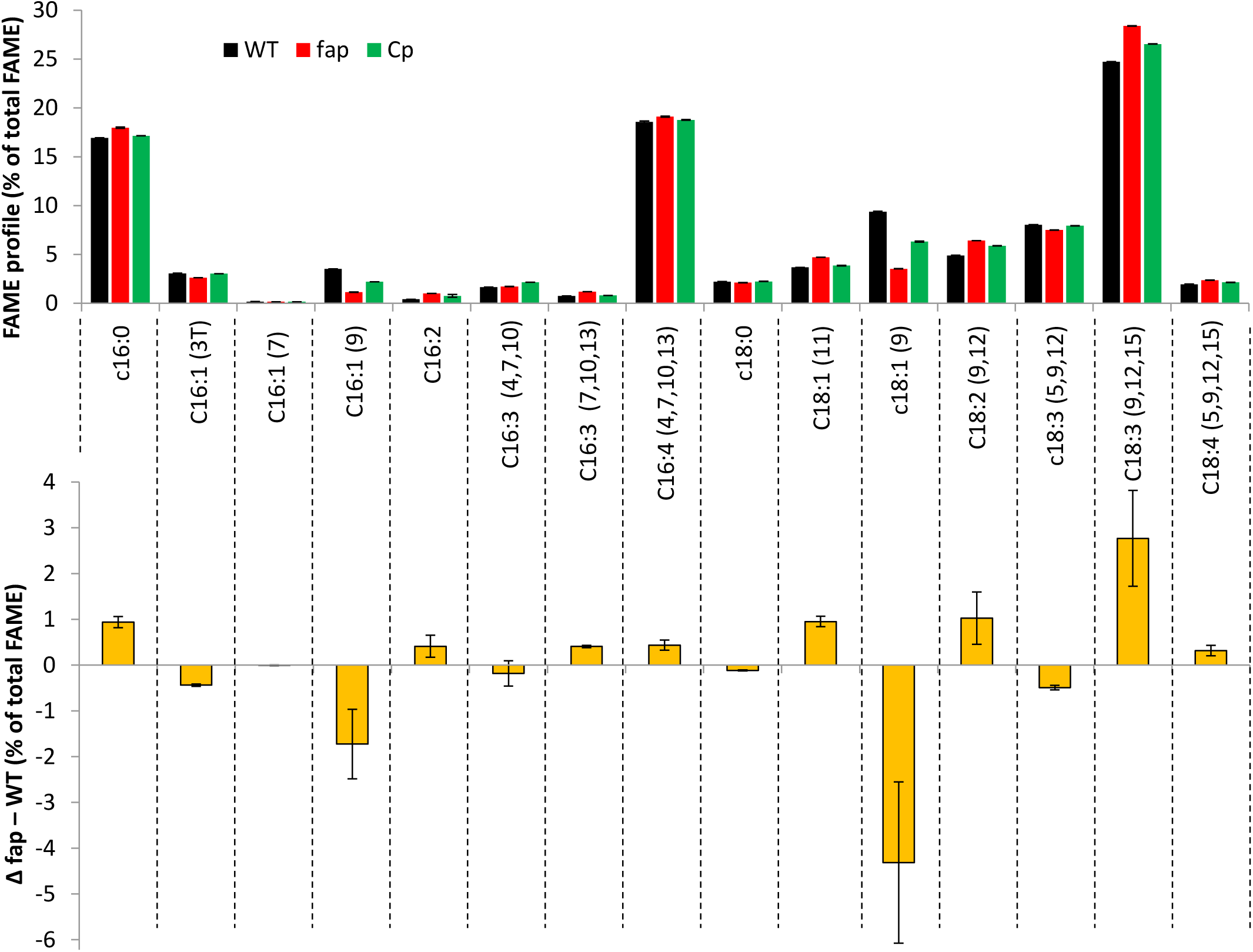
Fatty acid profile in mixotrophic conditions. Relative abundance of fatty acids methyl esters from transmethylation of whole cells analysed by GC-MS and expressed as percentage of total FAMEs. Cells grown in TAP medium, under 80 µmol photon m−2 s −1) in erlens (means ± SD; n = 3 biological replicates).

**Figure S10.**
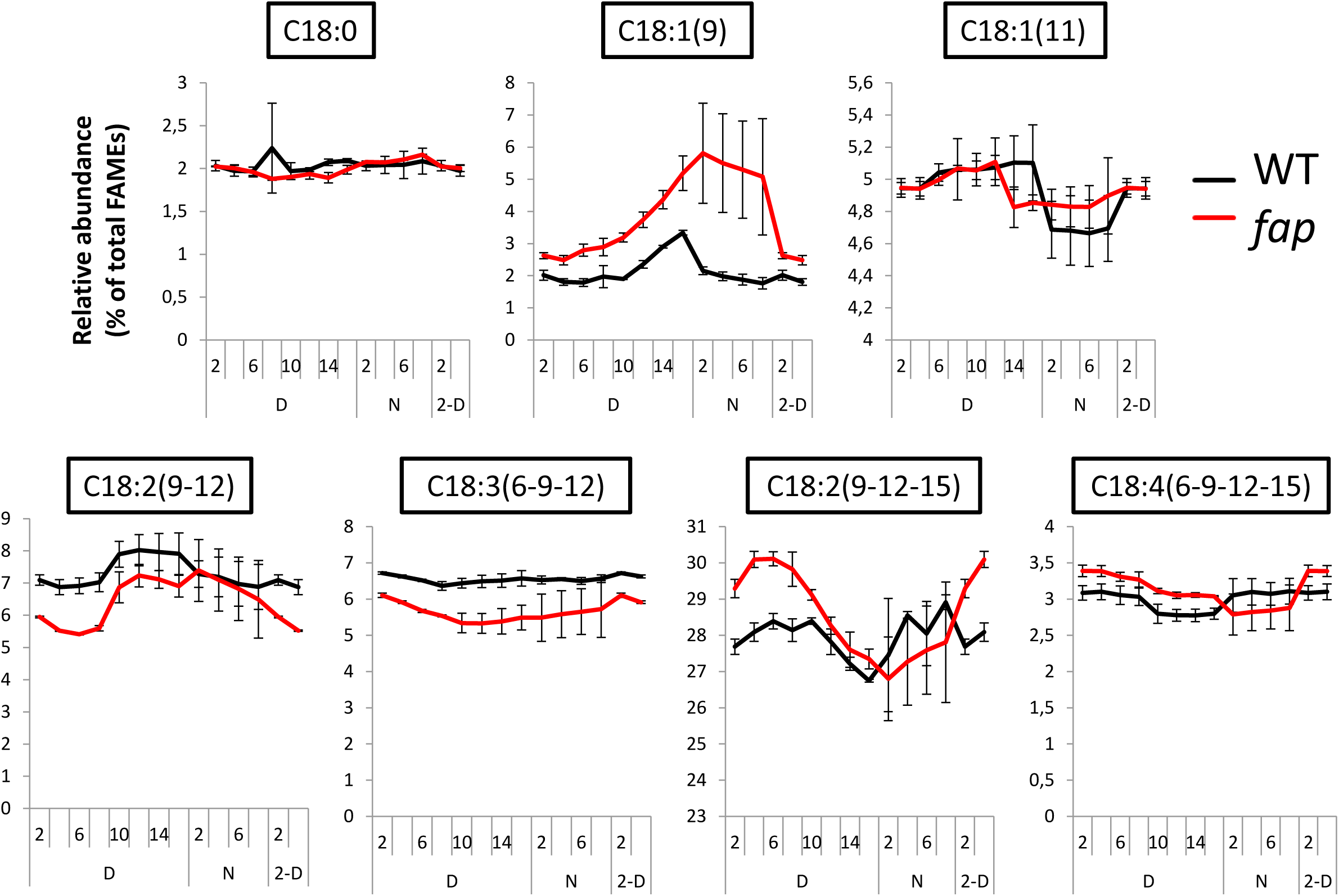
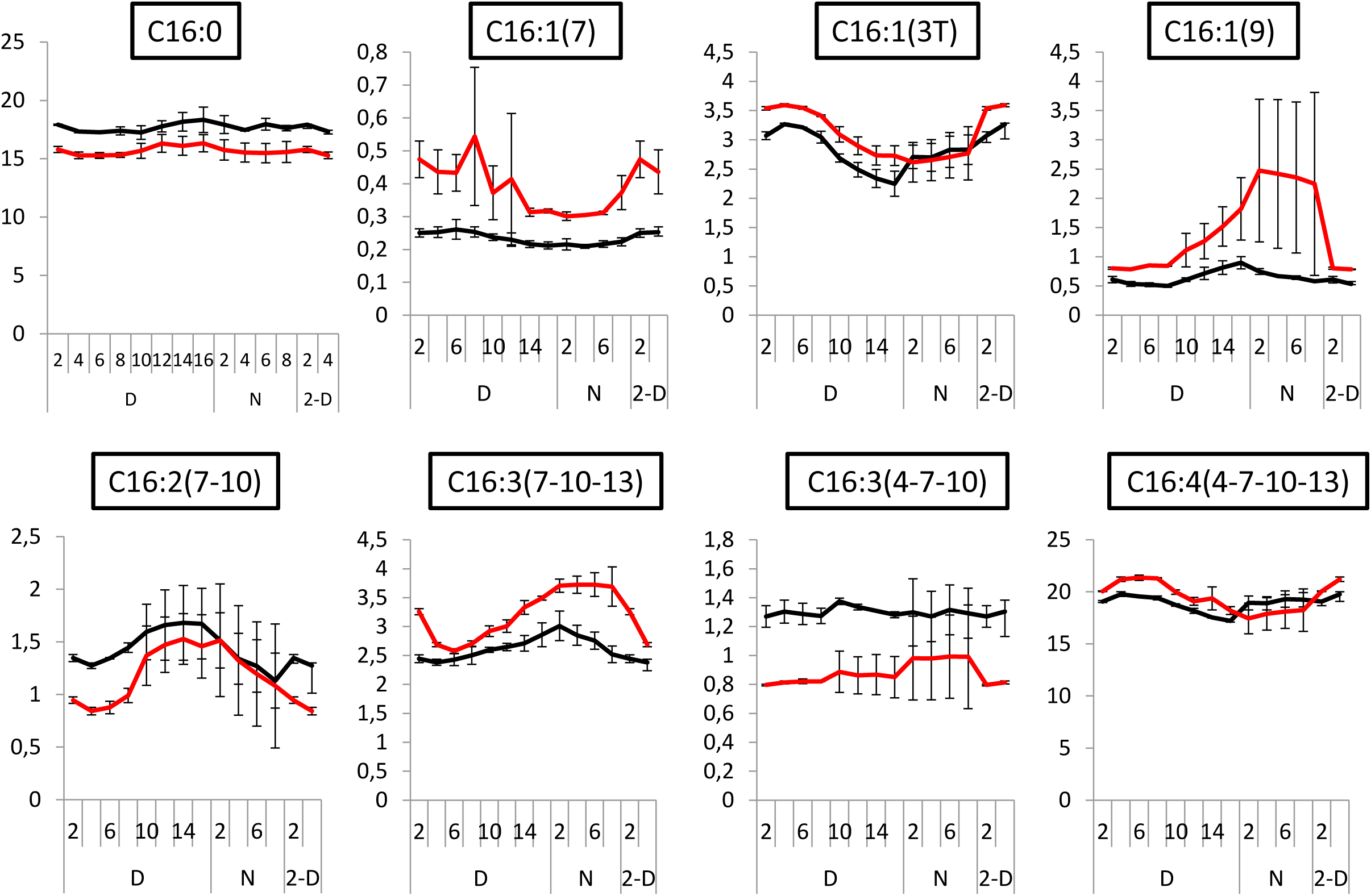
Variation in the proportion of each fatty acid in total fatty acids during cell cycle. Relative abundance of fatty acids methyl esters from transmethylation of synchronised cells analysed by GC-MS and expressed as percentage of total FAMEs along a day-night cycle, D : day (16 hours), N : night (8 hours), 2-D : first 2 measurements of the day to visualise the cycle (means ± SD; n = 3 biological replicates).

**Figure S11.**
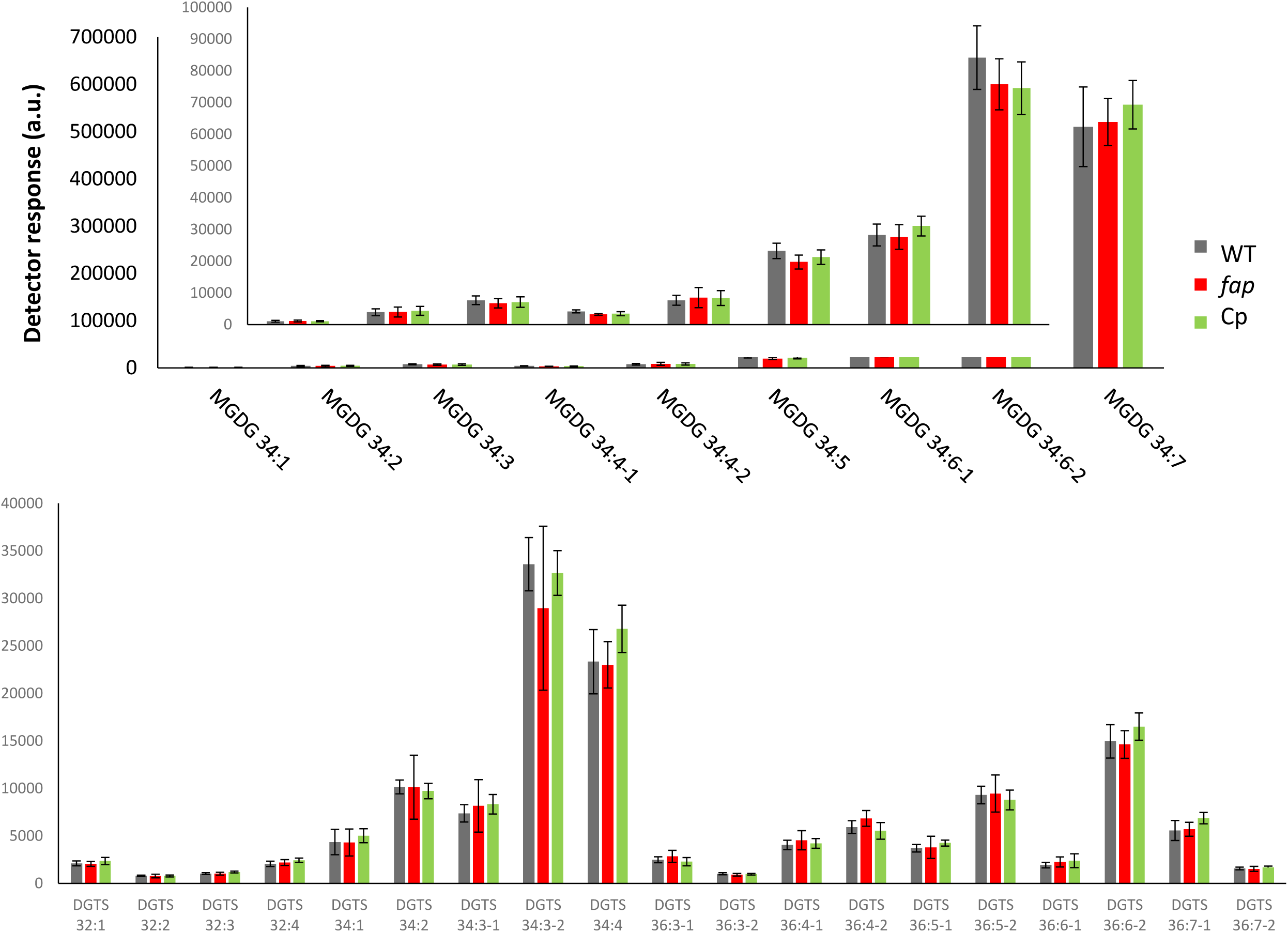
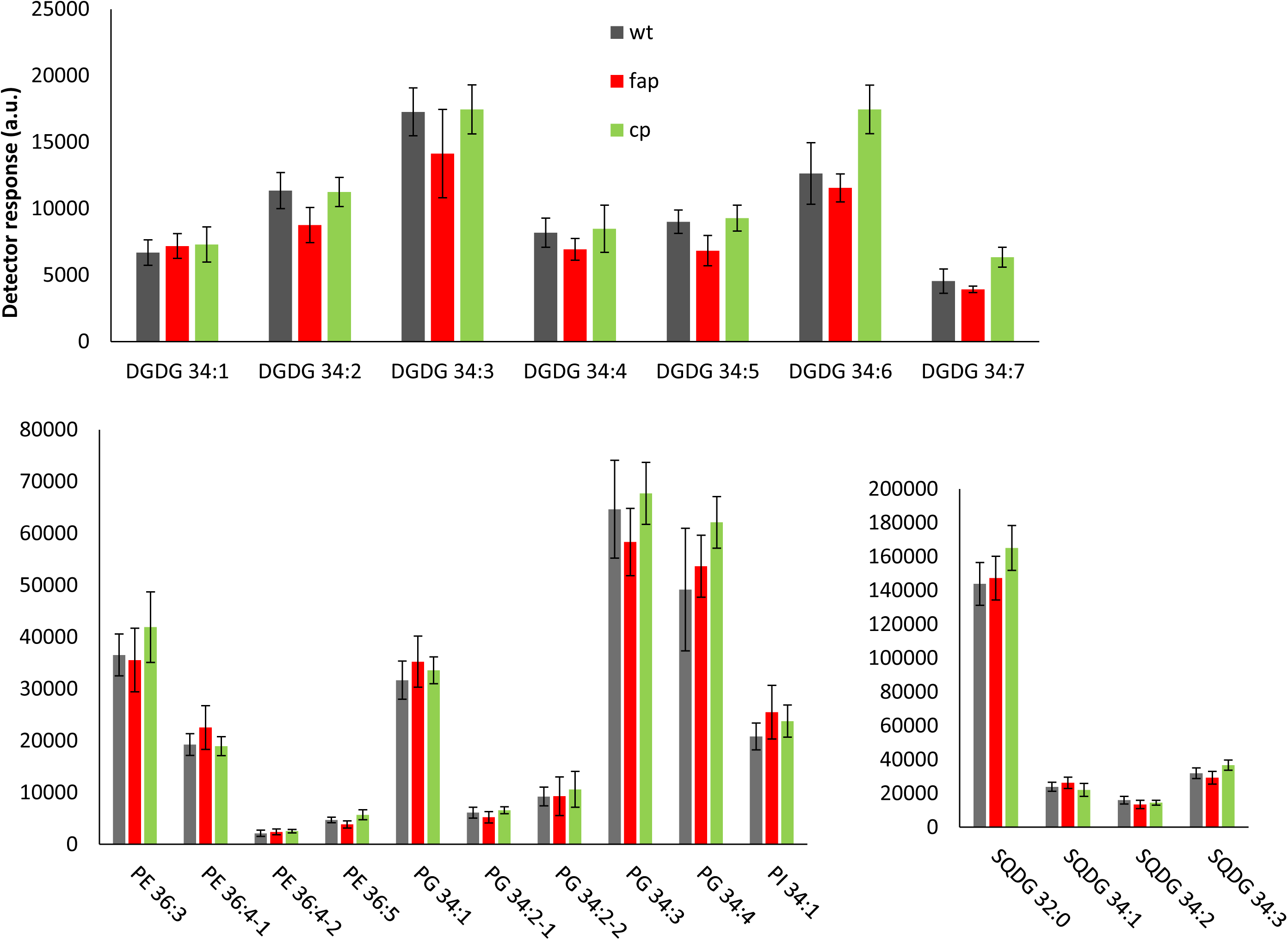
Profiles of major lipids in WT, *fap* and complementant strains. Relative abundance of lipids from LC-MS/MS analysis of whole cells on a total fatty acid basis. Species showing significant differences between WT and *fap* or Cp strains are shown in figure 7. Cells were grown in TAP medium, under 80 µmol photon m−2 s −1) in erlens (means ± SD; n = 9 biological replicates). On 2 pages

**Figure S12.**
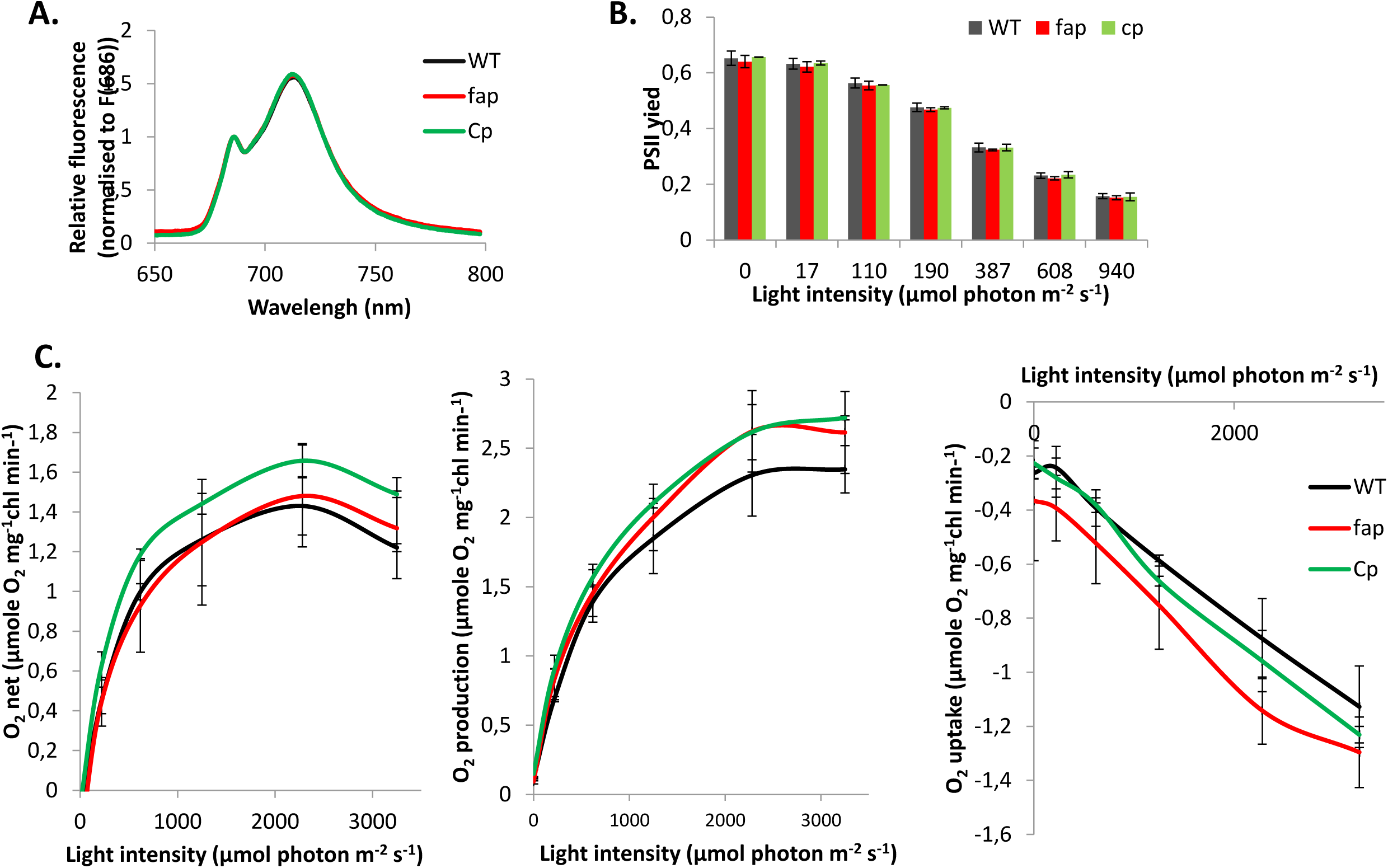
Photosynthetic activity in fap and WT strains. A, 77K fluorescence spectrum. B, Photosystem II operating yield under various light intensities. C, Oxygen uptake and production measured by membrane inlet mass spectrometry after acclimation for 2 minutes at each light intensity. Values are mean ± SD (n = 3 independent cultures)

**Figure S13.**
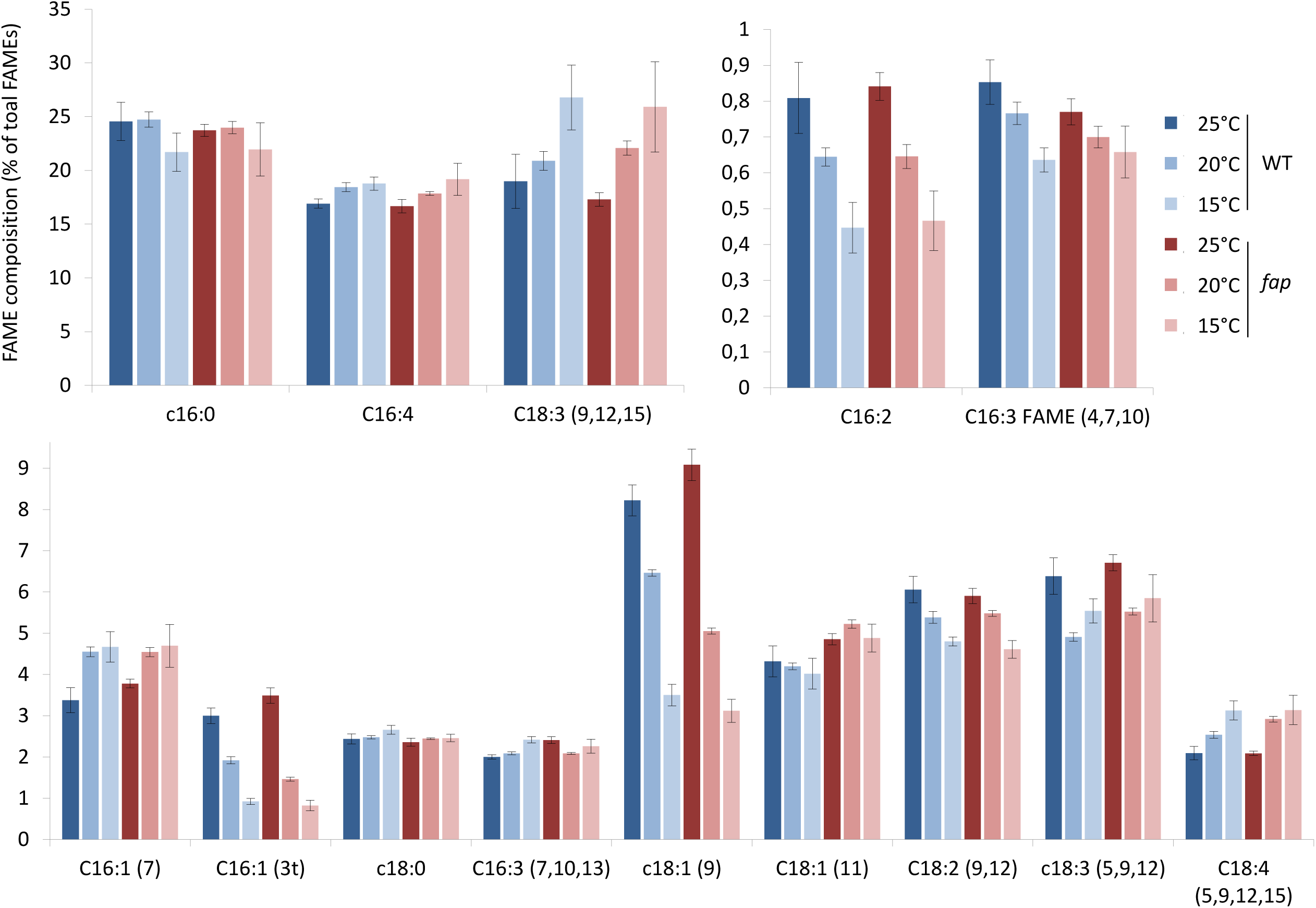
Fatty acid acclimation to cold conditions. Fatty acid content was analysed by GC-MS after transmehylation. Cells were grown in photobioreactor, in turbidostat mode, in TAP medium. (Mean of 4 replicates, error bar show standard deviation).

**Table S3.**
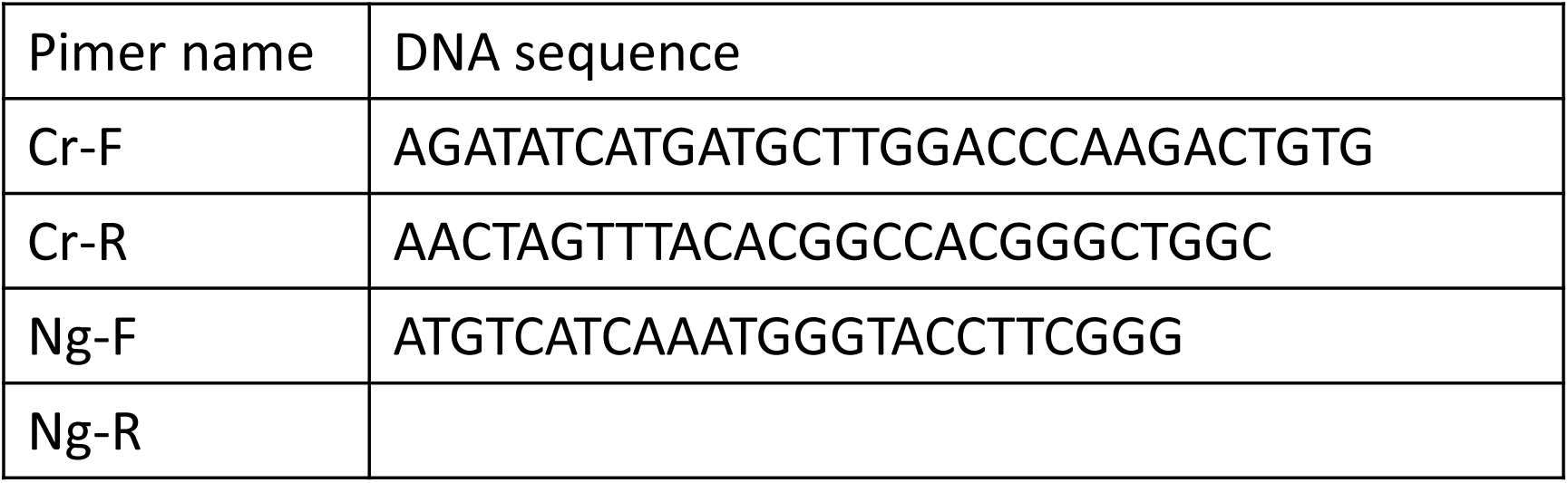
Primer used for PCR of FAP gene from cDNA of *C. reinhardtii* (Cr) and *N. gaditana* (Ng).

