## Supplementary figures and images for "Fatty acid photodecarboxylase is an ancient photoenzyme responsible for hydrocarbon formation in the thylakoid membranes of algae"

### Supplemental Figure 2

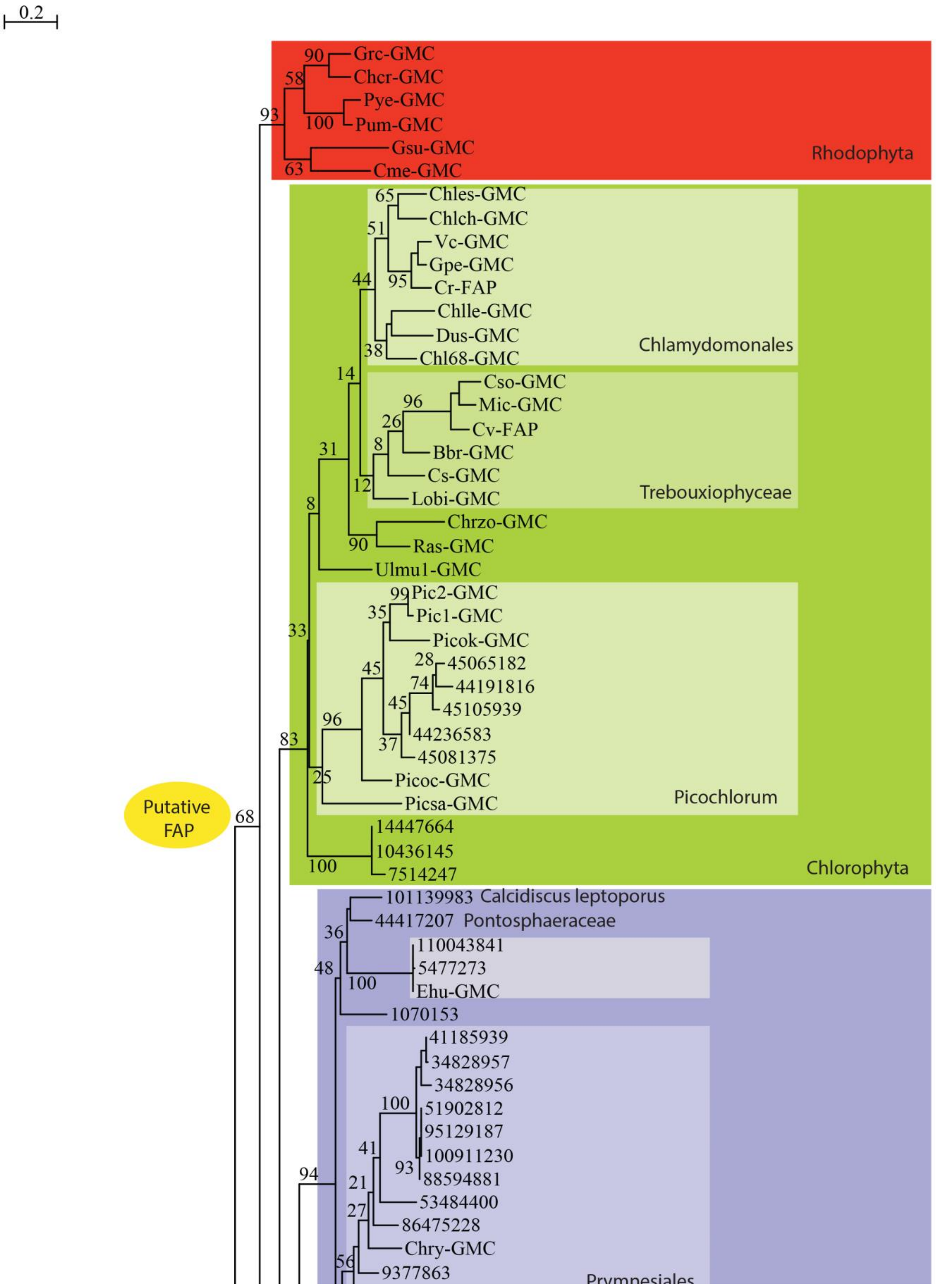

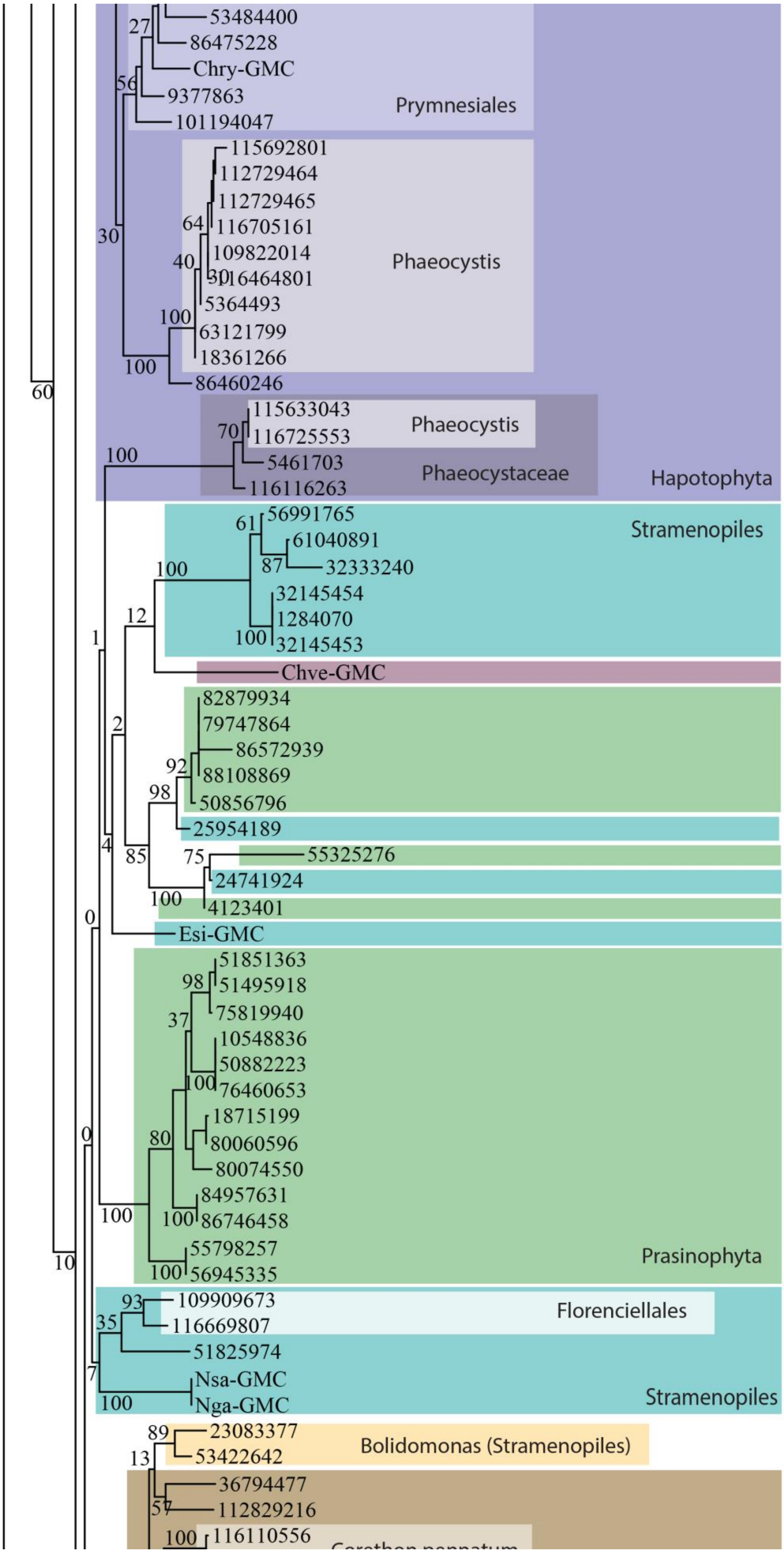

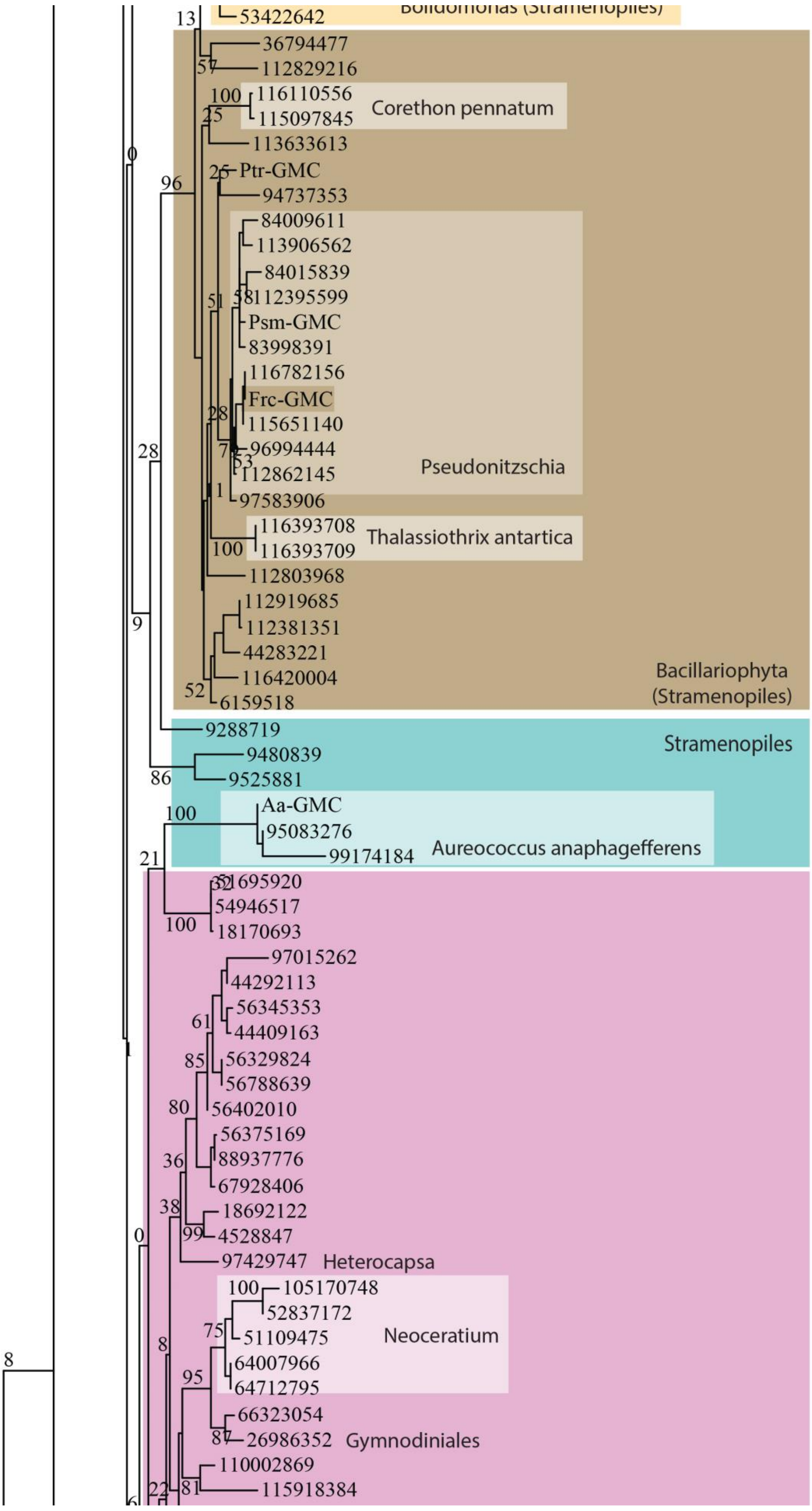

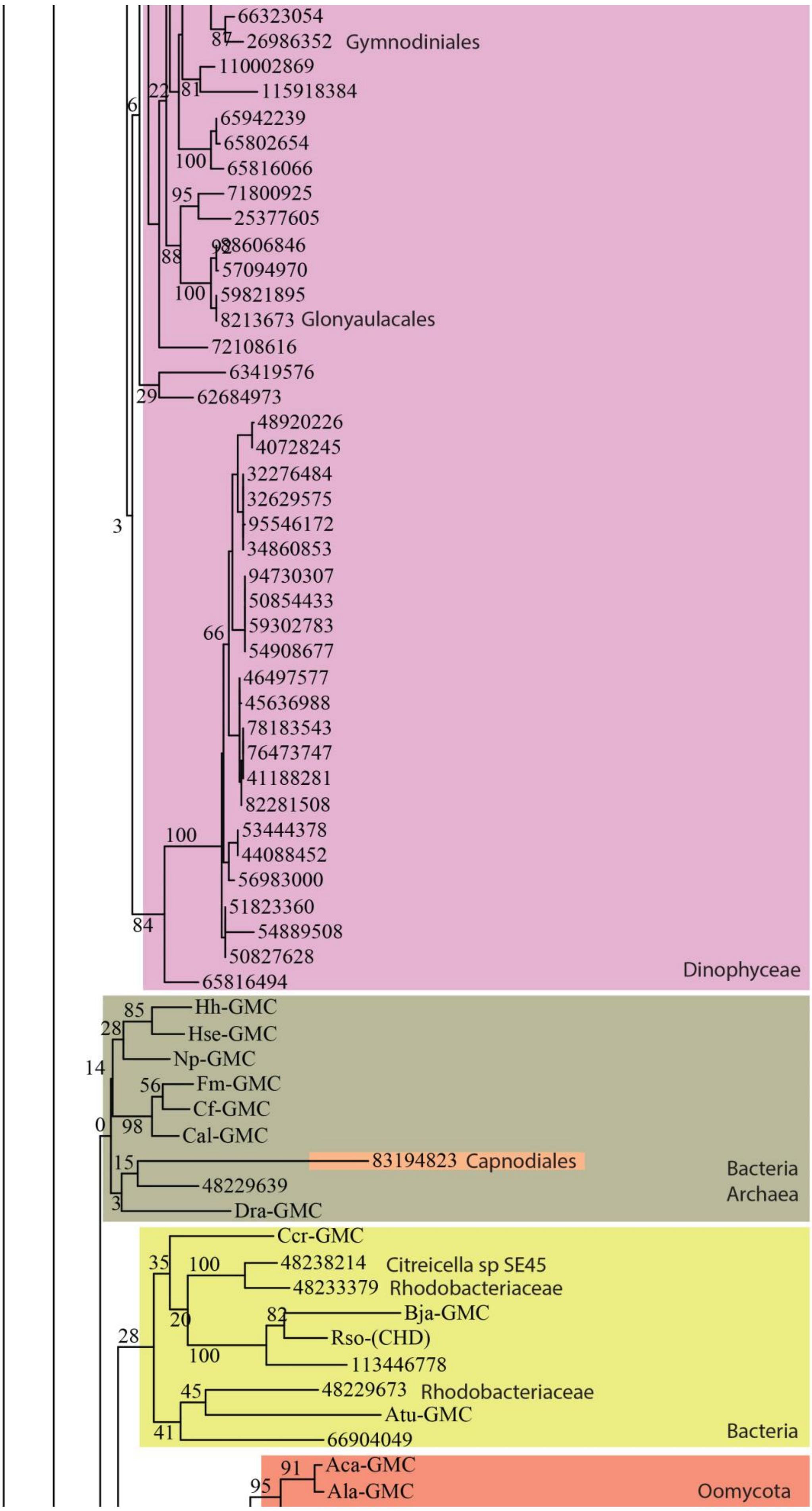

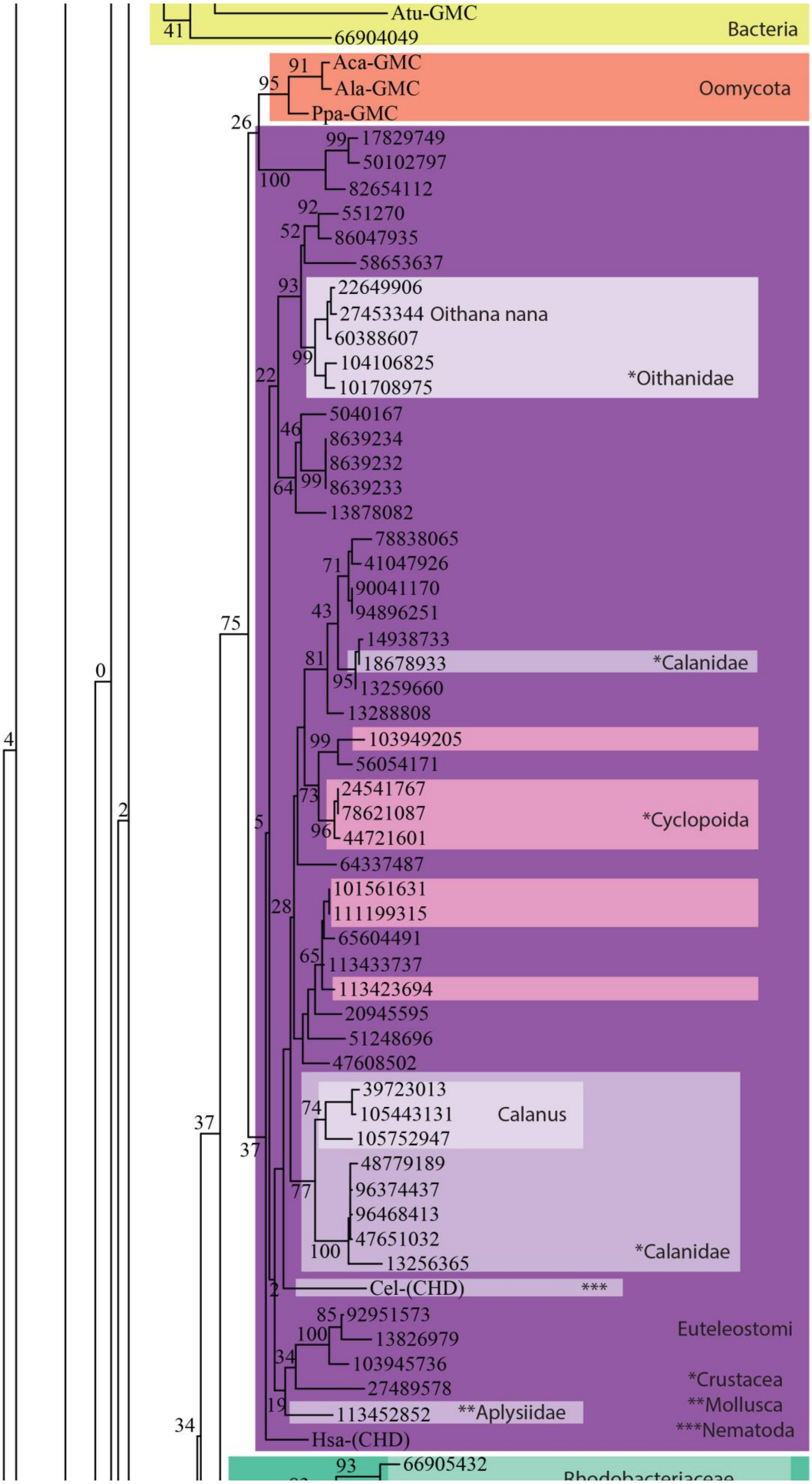

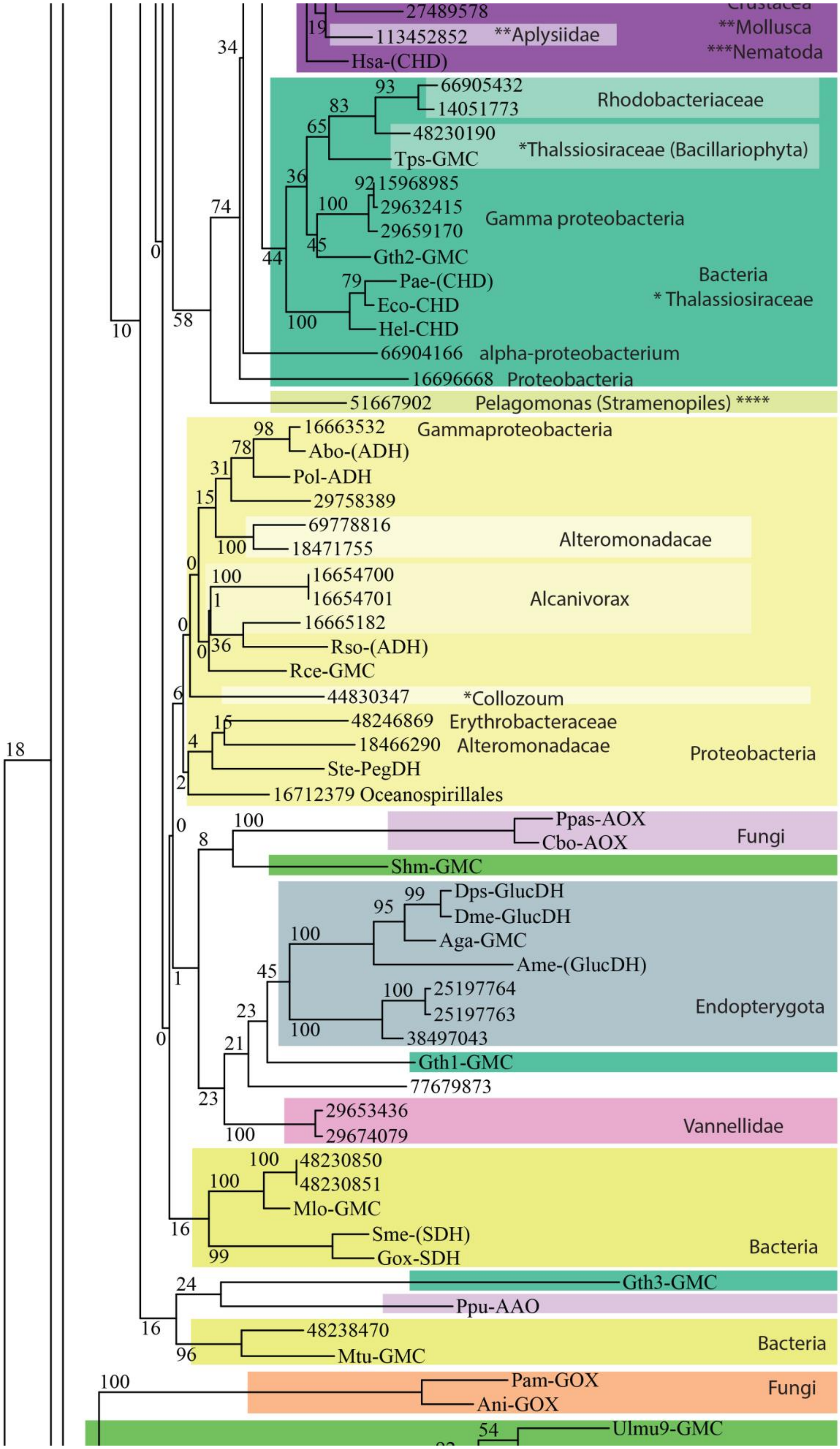

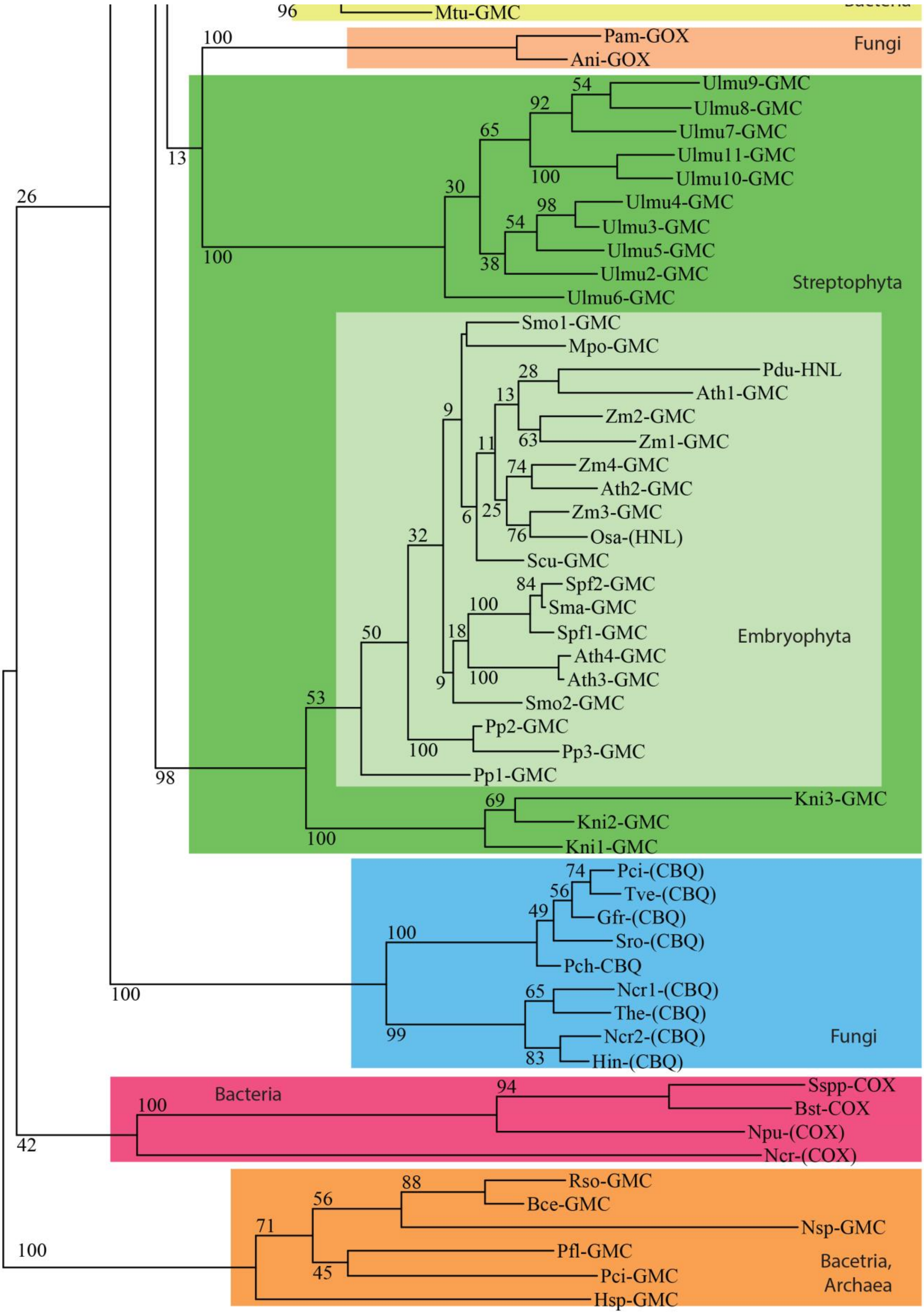
